# Evidence that host-mediated epigenetic modifications regulate gene expression in *Wolbachia pipientis*

**DOI:** 10.1101/2024.04.08.588587

**Authors:** Stella Papaleo, Simona Panelli, Ibrahim Bitar, Lodovico Sterzi, Riccardo Nodari, Francesco Comandatore

## Abstract

*Wolbachia pipientis* is an obligate intracellular bacterium, associated with several arthropods and filarial nematodes. *Wolbachia* establishes strict symbiotic relationships with its hosts, with the consequent loss of many genes and regulatory regions. Despite this, experimental studies show that *Wolbachia* gene expression is coordinated to host needs, but the mechanism is still unknown. The first published RNA-Seq study on *Wolbachia* evidenced a strong differential expression of a DNA methyltransferase (MTase). In bacteria, this enzyme methylates either adenines or cytosines on specific motifs, contributing to the regulation of gene expression. In this work, we tested the hypothesis that the activity of MTase modulates the expression of *Wolbachia* genes. We first determined the methylation motif of the *Wolbachia* MTase by expressing it in *Escherichia coli*. Surprisingly, the experiment revealed that the *Wolbachia* MTase methylates both adenine and cytosine, without recognising highly specific motifs. Then, re-analysing data from six RNA-Seq studies, we found that the nucleotide content of *Wolbachia* genes correlates with their expressions, with a pattern compatible to be a consequence of the DNA methylation. Lastly, we identified MTase as the *Wolbachia* gene with the most conserved binding site for the Ccka/CtrA signalling transduction system, a mechanism likely involved the host-bacterium communication. Overall, these findings suggest a cascade mechanism in which the host activates the *Wolbachia* Ccka/CtrA signalling system, thus inducing the expression of the MTase gene. Then, the subsequent DNA methylation will affect the expression of several *Wolbachia* genes on the basis of their cytosine and adenine content.

## Introduction

*Wolbachia pipientis* (from here *Wolbachia*) is an intracellular bacterium belonging to the *Alphaproteobacteria* class. It is the most prevalent endosymbiotic microbe in the animal world, colonising many arthropods species and filarial nematodes(1–4). In arthropods, the bacterium can manipulate the reproduction of the host(5), protect it from viruses(6, 7) and provide nutrients(8). In filarial nematodes, genomic analyses and experimental evidence revealed that *Wolbachia* is essential for the host survival and development (9). The bacterium is strictly intracellular, with a tropism for microtubules and with the ability to colonise endoplasmic reticulum (ER), with important effects on host cell replication(2). It is mainly mother-to-offspring transmitted, being actively transported by specialised cells of the host from the soma to germ cells (2). Nevertheless, horizontal transfer among host individuals can occur (10, 11), ehnancing the bacterium spreading (11).

As for many other strictly host-associated bacteria, the *Wolbachia* genome underwent an important reduction during its evolution, reaching a size that ranges from ∼0.8 Mb to ∼1.8 Mb (BV-BRC representative genomes)(12). As observed for several endosymbiont bacterial species, genome reduction passes through the increasing of the number of mobile elements (e.g. Insertion Sequences) and the loss of many genes(13–15). During this process, the structure of the genome and the order of genes can change dramatically(14), leading to the disruption of several operons and loss of most gene promoters. Interestingly, RNA-seq studies have highlighted that *Wolbachia* gene expression can be influenced by host environmental/developmental conditions(16–22). This suggests that *Wolbachia* likely has a mechanism for regulating gene expression, despite the low number of promoters indicating it may not depend solely on transcription factors.

The first published *Wolbachia* RNA-seq study(16) compared gene expression of *Wolbachia* colonising somatic tissues and gonads in females and males of the filarial nematode *Onchocerca ochengi*. One of the strongest differential expressions was reported for a gene annotated as a S-adenosyl-methyonine-dependent (SAM)-dependent DNA methyltransferase (hereafter ”MTase”). In general, DNA methyltransferase enzymes are able to methylate either adenine or cytosine on specific motifs of bacterial DNA. The most studied MTase are the DNA adenine methyltransferase (Dam) and DNA cytosine methyltransferase (Dcm) of *Escherichia coli*. The first one methylates the adenine residues in the sequence GATC, while the other one targets the second cytosine within the motifs CCAGG and CCTGG (23, 24). Furthermore, in *Alphaproteobacteria*, one of most characterised methyltransferase is the CcrM, which methylates the adenine in the motif GAnTC, and it is involved in the regulation of the cell cycle in *Caulobacter crescentus* (*25*).

Despite DNA methylation has deeply been investigated in eukaryotes, as an epigenetic mechanism of gene expression, its role in prokaryotes is still poorly investigated and no studies have been published on *Wolbachia*.

Despite the effect of DNA methylation on gene expression has been deeply investigated in eukaryotes (i.e. epigenetics), its role in prokaryotes physiology is still poorly understood and no studies have been published on *Wolbachia.* At the state of the art, there is evidence that DNA methyltransferase enzymes are involved in fundamental bacterial processes, including proofreading during DNA replication, protection from foreign DNA and regulation of gene expression. A specific type of DNA methyltransferase, called orphan DNA methyltransferases, methylates gene promoters, changing their affinity for transcription factors with important effects on gene expression (23, 24).

In this study, we investigated the hypothesis that DNA methylation is a key mechanism for the regulation of the *Wolbachia* gene expression. First, we experimentally determined the methylation motif of the MTase enzyme encoded in the genome of the *Wolbachia* endosymbiont of *O. ochengi* (wOo). Then, we tested whether the frequency of this motif into gene sequences correlates with their expression, and whether the intensity of the phenomenon is related to the levels of expression of the MTase gene. Finally, we identified regulatory regions upstream of the MTase gene in *Wolbachia* genomes. Our results suggest that the host may specifically induce the expression of the *Wolbachia* MTase gene through a signalling transduction system located on the bacterium’s membrane. Subsequently, the MTase methylates the *Wolbachia* genome, altering the expression of methylated genes based on their nucleotide composition.

## Material and methods

### Experimental determination of the wOo SAM-dependent DNA methyltransferase methylation pattern

The wOo SAM-dependent DNA methyltransferase (MTase) was artificially synthesised with codon usage adaptation and cloned into the XhoI and SalI sites of the pCOLD III expression vector (Takara Bio). The recombinant construct was transformed into *Escherichia coli* StellarTM Competent Cells (*dam*–/*dcm*–) (Takara Bio) according to the supplier’s protocol (ClonTech; Protocol-at-a-Glance, PT5056-2). The *E. coli* strains were cultured in LB broth medium supplemented with ampicillin. Before inducing expression, PCR with custom primers targeting the gene of interest was performed on bacterial crude extracts to verify the outcome of transformation. MTase expression was then induced at 15°C, according to the protocol provided by the manufacturer for the Cold Shock Expression System of pColdTM plasmids (Takara Bio). 24 hours after the induction, the expression of the wOo MTase gene was verified by SDS-PAGE. DNA was then extracted using the NucleoSpin Microbial DNA Mini kit (Macherey Nagel) and subsequently subjected to long reads sequencing using Sequel I (Pacific biosciences). Assembly of reads was done using the Microbial Assembly pipeline provided by SMRT Link v.10.1 with a minimum seed coverage of 30x. The assembled genome was used as a reference in the downstream analysis. The methylation status of each base of the obtained reads was then determined using the ‘Base modification

Analysis’ pipeline included in SMRT-Portal. The pipeline performs base modification and modified base motifs detection. A control experiment was carried out transforming *E. coli* Stellar TM Competent Cells with a pCOLD III vector without the wOo MTase gene inserted. The 95th percentile of Mean QV and coverage values obtained from the control experiment were set as minimum thresholds for the identification of methylated bases on the transformed *E. coli* strain. These last analyses were performed using R.

### Dataset reconstruction and data normalisation

The table S2 and gff map files of the supplementary information of Chung et al. 2020 were retrieved (26). The table contains the gene expression quantifications computed by Chung and colleagues (2020) re-analysing seven RNA-Seq studies on *Wolbachia* (16–22), for a total of 128 samples (replicates) from 62 host conditions. More in details: i) four samples from two conditions on *Wolbachia* endosymbiont of *Onchocerca ochengi* (wOo) from Darby et al. 2012(16); ii) five samples from five conditions on *Wolbachia* endosymbiont of *Dirofilaria immitis* (wDi) from Luck et al., 2014(17); iii) six samples from two conditions on *Wolbachia* endosymbiont of *Drosophila melanogaster* (wMel), from Darby et al., 2014(18); iv) 12 samples from seven conditions on wDi from Luck et al., 2015(19); v) 55 samples from 24 conditions on wMel, from Gutzwiller et al., 2015(20); vi) 14 samples from seven conditions on *Wolbachia* endosymbiont of *Brugia malayi* (wBm) from Grote et al., 2017(21); vii) 32 samples from 15 conditions on wBm from Chung et al., 2019(22). Preliminarily, we excluded from the dataset all the samples from Luck et al., 2015(19) because only 8% of the gene expression quantifications were greater than 0. Thus, gene expression analysis was performed on a total of 116 samples from 55 host conditions, two for wOo, five for wDi, 22 for wBm and 26 for wMel.

Gene expression values were normalised to make them comparable among studies and conditions. For each condition in each study the gene expression values were expressed as

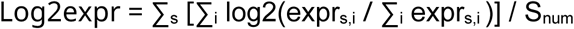

Where:

s = sample (or replicate); i = gene; exprs,i = the gene expression value of the gene i in the sample (replicate) s; Snum = total number of samples. This normalisation formula derives from the log2 fold change formula often used to compare two conditions in RNA-Seq studies:

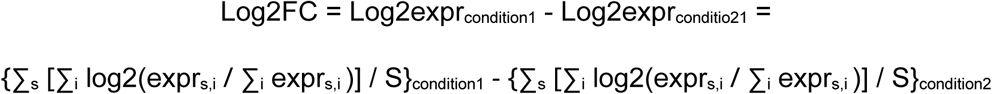

Log2FC allows us to compare values from different conditions in the same experiment. To make the normalised values more comparable among the studies, they were converted in percentiles. Indeed, the percentile may be less biassed by sample specific features, such as the number of genes.

### Genes composition parameters determination

The genome assemblies of wBm (AE017321.1), wDi (www.nematodes.org: wDi 2.2), wOo (HE660029.1) and wMel (AE017196.1) were retrieved and passed to Prodigal (27) for Open Reading Frame (ORF) calling. For each ORF, Codon Bias Index (CBI) (which measure how much a gene uses a subset of optimal codons, ranging from 0 for random using, to 1 for maximum bias), the effective number of codons (Nc) and the ORF length were computed using CodonW tool (Peden J.F. Analysis of codon usage. 1999. PhD Thesis, University of Nottingham, UK), while %A, %T, %G, %C were computed using an in-house Perl script. Conventionally, ORF sequences correspond to the transcribed messenger RNA (mRNA) sequence, which is complementary to the DNA template strand. Thus, the nucleotide composition of the genes is derived by complementing the ORF compositions: the %A of a gene is determined as the %T of the corresponding ORF, and so on. Finally, the %C/%A ratio is calculated. All subsequent analyses will focus on the nucleotide composition of the genes rather than on ORF compositions.

### Statistical analyses

The relationship between each gene composition parameter (CBI, Nc, length, %A, %T, %G, %C and C/A ratio, see above) and gene expression was investigated. More in detail, for each of the 55 samples included in the study, linear regression was used to assess the correlation between the global gene expression reported for that sample and each of the gene composition parameters. For each regression line, the slope and the p value were obtained. Slope values indicate the strength of change in gene expression in relation to the gene composition parameter: the higher the slope, the more the gene expression varies as a consequence of that gene composition parameter. In a second analysis, to investigate if MTase activity affects the expression of genes on the basis of their composition parameters, the correlation between the linear regression slope and the MTase gene expression was computed for each gene composition parameter.

The presence of other genes with expression patterns comparable to that of MTase gene was tested as follows. Exploiting the gene orthologous information included in the Table S6 of Chung et al., 2020 (26), we performed a Spearman co-correlation among the expressions of the 546 single copy core genes shared among wOo, wDi, wBm and wMel. The expression patterns were also investigated by Principal Component Analysis (PCA), using R.

### Comparison between species tree and MTase tree

A dataset including 112 high quality *Wolbachia* genome assemblies spanning the host genetic diversity was reconstructed, including 111 assemblies retrieved from the Genome Taxonomy DataBase (GTDB) and the wDi genome used for the previous analysis (indeed, the wBm, wMel and wOo assemblies were already present in the GTDB database) (see Table S1). The genome assemblies were subjected to Prodigal (27) for ORF calling, and the amino acid sequences were passed to OrthoFinder (28) for orthologue analysis. The nucleotide sequences of the single copy core genes were then retrieved, aligned using the MUSCLE tool (29), tested for recombinations using the PhiPack tool (30), trimmed using the trimal tool (-gt 0.5 setting) (31), and finally concatenated. The obtained concatenate was passed to RAxML8 (32) for phylogenetic analysis using the GTR+I+G model, as previously determined using ModelTest-NG tool (33). The orthologue group containing the MTase genes was determined and subjected to phylogenetic analysis following the same flow described above (using the TVM+I+G model and excluding the step of the concatenation of the genes). The obtained trees were then compared using the Cophylo R library (34).

### Analysis of the region upstream the methyltransferase gene

The positions and orientation of the MTase genes on the 112 genome assemblies were determined by BLASTn searches. Then, for each genome assembly, the 100 nucleotides upstream of the MTase gene were extracted and screened for the presence of some of the most important regulatory regions: i) CtrA binding site (motif TTAA-N7-TTAA (35)); ii) Pribnow box (motif TATAAT (36)); iii) CAAT box (motif HYYRRCCAWWSR(37)); iv) GC box (motif WRDRGGHRKDKYYK(37)); v) TATA box (motif TATAWAWR(37)). Perfectly matching motifs and one-mismatched motifs were considered for further analyses. The presence of *cck*A and *ctr*A genes in the 112 *Wolbachia* genomes was then evaluated by BlastP search, using as reference the *cck*A and *ctr*A gene sequences already reported in Lyndsey 2020 (38). Finally, the positions of the canonical regulatory regions upstream the MTase gene, and the presence/absence of *cck*A/*ctr*A genes in the 112 *Wolbachia* genomes were visualised using the gplots R library.

### Analysis of the regions upstream all genes in the 112 *Wolbachia* genomes

On the basis of previous results on the MTase gene, TATA boxes were searched between positions -20 and -70 upstream of all the genes of the 112 *Wolbachia* genomes, and CrtA binding sites between the positions 0 and -30. The presence of short AT-rich sequences, like TATA box and CtrA binding site, in AT-rich genomes (such as *Wolbachia*) could be due to the chance. The possible functionality of these sequences has been assessed by investigating whether TATA or CtrA binding sites are enriched upstream of specific genes in the *Wolbachia* genomes. More in detail, for each of the ortholog genes previously identified using OrthoFinder (see above), the frequency of *Wolbachia* strains having upstream TATA or CtrA binding site was investigated using R. Lastly, both for TATA box and CtrA binding site, the genes present in at least 100 out of 112 *Wolbachia* genomes and having the regulatory box in >80% of the genomes were retrieved and annotated using the Clusters of Orthologous Groups (COG) database.

### Analysis of the genes with high and low G/T ratio

One of the gene composition parameters found to be more correlated to gene expression and MTase action was C/A ratio. Thus, the expression of the genes with higher or lower C/A ratio is expected to be more affected by the action of the MTase enzyme. To investigate the effects of this mechanism on the *Wolbachia* metabolism and physiology, genes with low or high C/A ratio values were studied as follows. All the amino acid sequences from the 112 *Wolbachia* strains included in the study were annotated on the basis of the Clusters of Orthologous Genes (COG) database. For each strain, among the COG-annotated genes, the 10 with lower C/A ratio (“low C/A ratio genes”) and the 10 with higher C/A ratio (“high C/A ratio genes”) were retrieved. The pattern of presence/absence and C/A ratio value of all these retrieved genes was graphically investigated by producing a heatmap using the gplots R library.

## Results

### Experimental determination of the wOo SAM-dependent DNA methyltransferase methylation pattern

The expression of the wOo MTase into the Stellar *E. coli dam*-/*dcm*- strain followed by the SMRT PacBio sequencing allowed to identify 166 high quality methylated bases: 151 on cytosine (classified as m4C - N4-methylcytosine) and 15 on adenine (classified as m6A - N6- methyladenine) (see Table S2 and Figure S1). Investigation of the methylation pattern revealed that m4c methylations occurred in a pattern that was not well conserved, whereas m6A methylation involved a guanine-rich region at positions -4, -8 and -19 (Figure 1).

**Figure 1.**
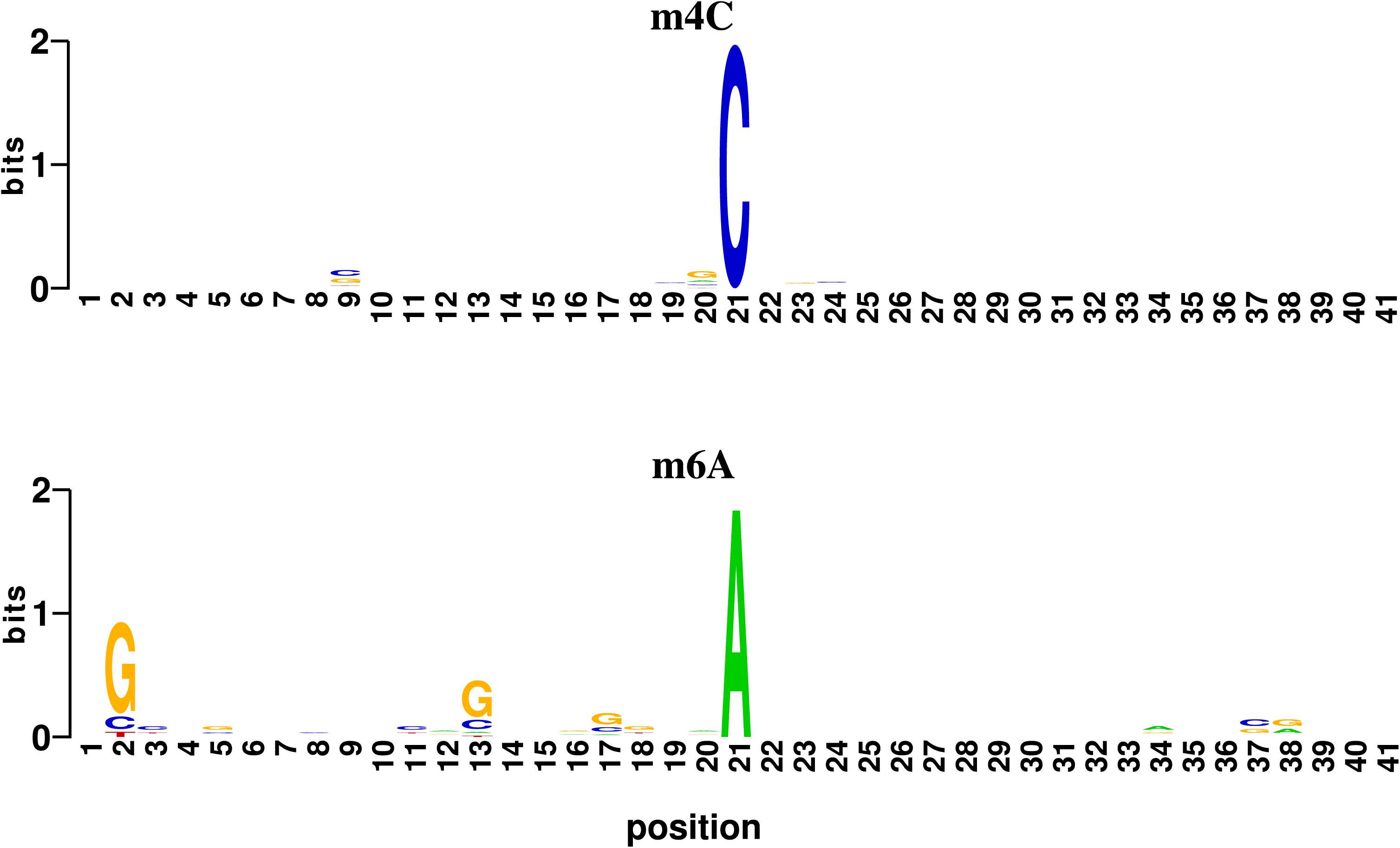
Methylation patterns of the DNA methyltransferase gene of *Wolbachia* endosymbiont of *Onchocerca ochengi* (wOo) Sequence logo of the sequences flanking (from -20 to +20 positions) the methylated positions found in the genome of Stellar *Escherichia coli* dam-/dcm- strain expressing the DNA methyltransferase gene of *Wolbachia* endosymbiont of *Onchocerca ochengi* (wOo). The height of the letters depends on the conservation of the position among the sequences. a) the sequence logo of the m4C methylations on cytosines; b) the m6A methylation of adenine.

### Dataset reconstruction

For each of the six datasets included in the study, the normalised gene expression values were obtained as described in material and methods. The ORF sequences of the wOo, wDi, wBm and wMel genome assemblies were analysed to determine the CBI, Nc, length, %A, %T, %G, %C and %C/%A (from here C/A ratio). The obtained information about gene composition parameters was merged to relative gene expression data (Supplementary information available on repository…).

### Statistical analyses

The slope and p value of the linear regression analyses performed between the eight gene composition parameters (CBI, Nc, gene length, %A, %T, %G, %C and C/A ratio) and the gene expression data obtained for each of the 55 samples are reported in Table S3. Length and %C significantly correlate with gene expression in all the samples included in the study (100%), while %T in 53/55 (96%), CBI 42/55 (78%), %A 40/55 (73%), C/A ratio 31/55 (56%), %G 22/55 (40%) and Nc 7/55 (13%). The obtained slopes for the regression lines range as follows: for gene length from 0.009 to 0.029, %C from 0.79 to 5.3, CBI from 30.9 to 71.7, %A from -1.8 to 1.1, C/A ratio from 18.5 to 90.6, %G from -1.3 to 2.2 and Nc from 0.2 to 0.8. Subsequently, for each of the gene composition parameters (and for each of the 55 samples) the correlation between the levels of expression of the MTase gene and the regression slope obtained above (indicative of the strength of the correlation between the gene expression data of the sample and the gene composition parameter) was tested by linear regression. The aim was to investigate if MTase activity affects the expression of genes on the basis of their composition parameters. The obtained correlations were significant for all parameters except %G, which had a p-value of 0.08. For %G, %C, %T, C/A ratio, Nc, and length, the slope increases with higher DNA MTase expression. Conversely, %A and CBI exhibit a decreasing trend with increasing of the MTase expression (Figures 2-3 and Figures S2-S7).

**Figure 2.**
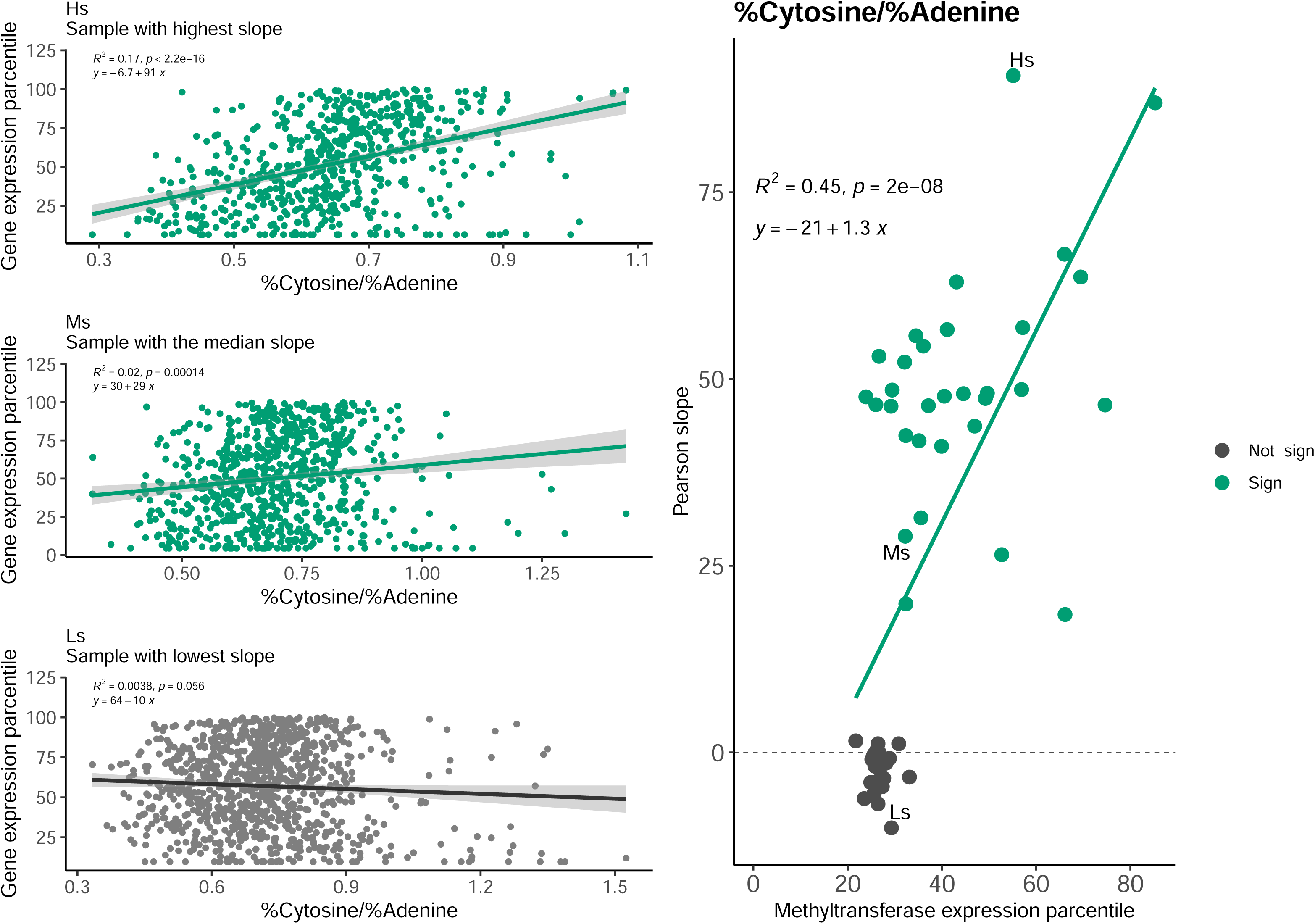
Correlation between the expression of genes and their %C / %A ratio For each of the 55 samples included in the study, the correlation between the expression of the genes and the relative %C/%A (“C/A ratio”) was investigated. The scatter plots of C/A ratio vs gene expression of three exemplificative samples selected from the 55 samples are reported: a) the correlation plot with highest slope (Hs), b) the correlation plot with the medium slope (Ms) and c) the correlation plot with the lowest slope (Ls). In each of these three plots, colour is green whether the correlation is significant (p value < 0.05) and grey if it is not, the regression line is coloured in dark grey and the statistics are reported on the chart. In figure d, the correlation plot between the slopes obtained for each of the 55 samples, and the relative expression of the DNA methyltransferase gene is reported. Points relative to significant correlations (p value < 0.05) are in green while non significant in grey. The regression line is in green and the statistics are reported on the chart.

**Figure 3.**
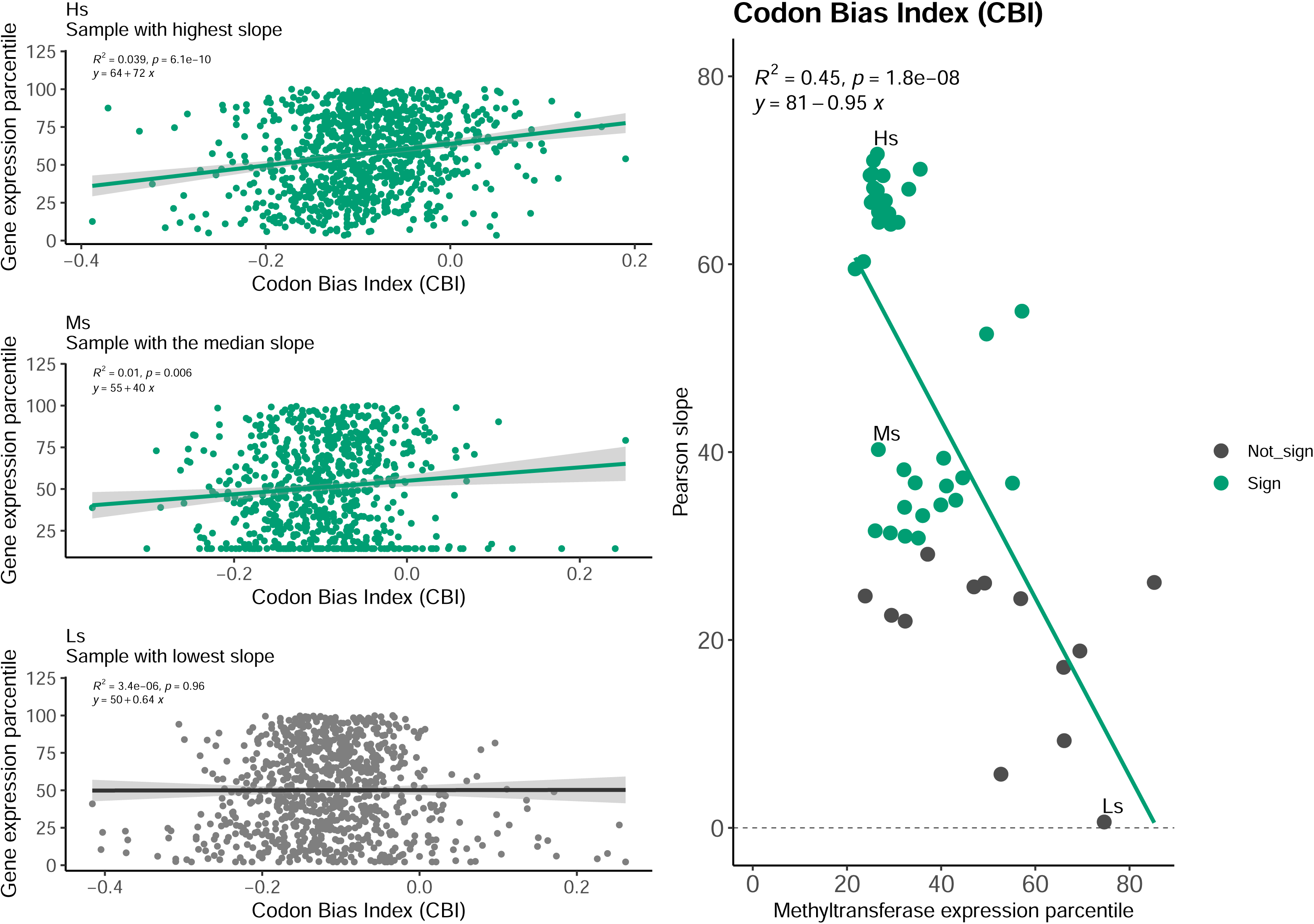
Correlation between the expression of genes and their Codon Bias Index (CBI) For each of the 55 samples included in the study, the correlation between the expression of the genes and the relative Codon Bias Index (CBI) was investigated. The scatter plots of CBI vs gene expression of three exemplificative samples selected from the 55 samples are reported: a) the correlation plot with highest slope (Hs), b) the correlation plot with the medium slope (Ms) and c) the correlation plot with the lowest slope (Ls). In each of these three plots, colour is green whether the correlation is significant (p value < 0.05) and grey if it is not, the regression line is coloured in dark grey and the statistics are reported on the chart. In figure d, the correlation plot between the slopes obtained for each of the 55 samples, and the relative expression of the DNA methyltransferase gene is reported. Points relative to significant correlations (p value < 0.05) are in green while non significant in grey. The regression line is in green and the statistics are reported on the chart.

Co-correlation analysis among the expression patterns of the single copy core genes shared among wOo, wDi, wBm and wMel showed that no other gene has an expression pattern overlapping that of the DNA MTase gene. Indeed, the highest correlation value was 0.84 and the median value was 0.02 (Figure S8). In general, the absence of clusters of co-expressed genes is also evident from PCA analysis (Figure S9).

### Comparison between species tree and MTase tree

Orthology analysis identified a total of 3,561 orthologous groups, including 263 single copy core genes. None of the single copy core genes were recombined. After trimming of gene alignments, the obtained concatenate had a length of 223,833 bp. On the other side, the MTase gene alignment was 1,342 bp long. The species tree obtained from the 263 single copy genes and the MTgene tree are reported in Figure S10. The obtained topologies were mainly congruent, with few signs of Horizontal Gene Transfer (HGT) events.

### Analysis of the region upstream the methyltransferase gene

The MTase gene was found in all the 112 genome assemblies included in the study. Unfortunately, for one genome assembly (accession GCF_018454445.1, *Wolbachia* endosymbiont of *Rhagoletis cerasi* - wCer5) the MTase gene was located on the extreme of a contig, and thus it was not possible to extract the 100 bp upstream of the gene transcription initiation site. The presence of CtrA binding site, Pribnow, CAAT, GC and TATA boxes upstream the MTase gene was assessed. Among these, at least one canonical sequence (i.e. perfect match) was found only for CtrA binding site, TATA and Pribnow boxes. In particular, 104 out of the 111 regions upstream of the MTase gene contained canonical TATA boxes (Figure S11), 100 out of 111 canonical CtrA binding sites (Figure S12) and five canonical Pribnow boxes (Figure S13).

### Analysis of the regions upstream all genes in the 112 *Wolbachia* genomes

The assessment of the presence of CtrA binding site and TATA boxes upstream of all the genes of the 112 *Wolbachia* genomes included in the study (see above) led to the discovery of a total of 2,210 CtrA binding sites and 12,283 TATA boxes. The CtrA binding site was predominantly found upstream of the MTase gene, but it was also frequently present upstream of the Formate hydrogenlyase subunit 3/Multisubunit Na+/H+ antiporter, which is involved in energy production (see Table S4 and Figure S14). Instead, the TATA boxes were found with high frequency upstream of 12 genes (see Table S4 and Figure S15), including the gene coding for the CtrA DNA binding response regulator.

### Analysis of the genes with high and low C/A ratio

A total of 57 low C/A ratio genes and 53 high C/A ratio genes were found. Among the low C/A ratio genes, the most frequent COG categories were “Translation, ribosomal structure and biogenesis” (J) with 13/57 (23%) genes, “Mobilome: prophages, transposons” (X) with 8/57 (14%), “Posttranslational modification, protein turnover, chaperones” (O) with 7/57 (12%) and “Transcription” (K) with 5/57 (9%) genes. Conversely, among the high C/A ratio genes, the most frequent COG categories were “Energy production and conversion” (C) with 15/53 (28%) genes, “Inorganic ion transport and metabolism” (P) with 11/53 (21%), “Lipid transport and metabolism” (I) with 5/53 (9%) genes. The presence/absence and C/A ratio patterns are shown in Figure 4.

**Figure 4.**
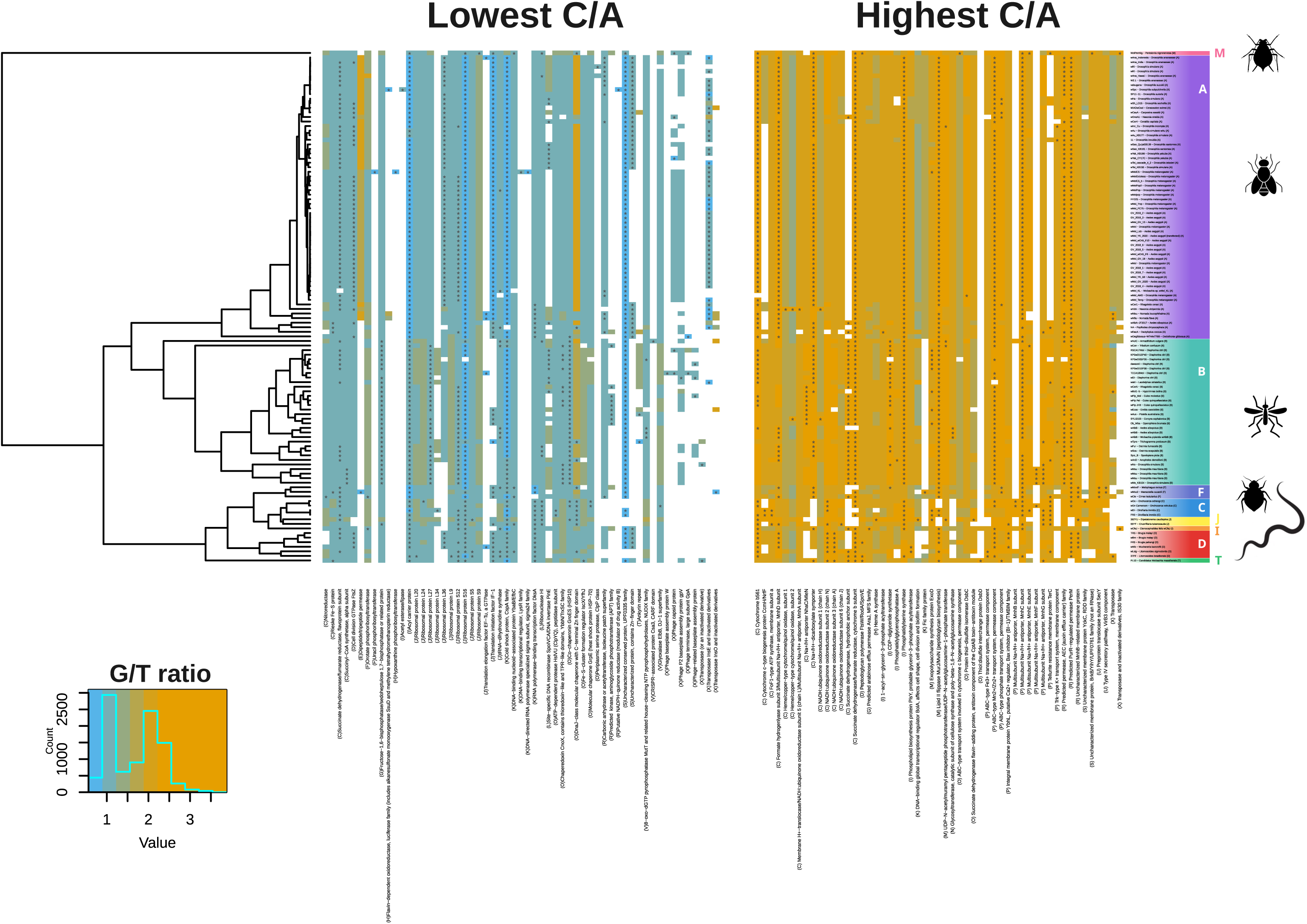
COG annotated genes with highest or lowest C/A ratio The heatmap shows the pattern of the genes with high or low C/A ratio among the 112 *Wolbachia* strains included in the study. On the left, the Maximum Likelihood phylogenetic tree obtained from concatenate of the single copy core genes. In the middle, heatmap reports the pattern of the 10 lower C/A ratio COG annotated genes: rows refer to strains (matching with the phylogenetic tree on the left) and columns to genes (COG annotation is reported). Cell colours report the C/A ratio value following the legend on lower left. Asterisks are reported for genes among those in the 10 lower C/A ratio for the relative strain. More on the right, a heatmap reports the 10 higher C/A ratio COG annotated genes. Finally, on the right, *Wolbachia* strains hosts and supergroups are shown.

## Discussion

*Wolbachia* is one of the most widespread endosymbiotic bacteria (10), with an important impact on the survival and/or reproduction of several arthropod and filarial nematode host species (2, 5, 39)(2, 3, 39). The close symbiotic relationship with the host has had important consequences on the bacterial genome structure, including frequent rearrangements (14) and the loss of many genes and the loss of regulatory regions (40), with the disruption of several operons and promoters. The loss of promoters could have limited evolutionary costs, considering that the intracellular niche of the bacterium protects it from external environment stresses. However, experimental studies showed the existence of *Wolbachia* genes differentially expressed in host samples from different body compartments, developmental stages and/or from hosts subjected to environmental conditions (26). Nevertheless, at the state of the art, the gene regulation mechanisms in *Wolbachia* are poorly understood.

One of the most differentially expressed genes between *Wolbachia* colonising the soma and gonads of the filarial nematode *Onchocerca ochengi* (wOo) was a SAM-dependent DNA methyltransferase (MTase) (16). Generally, methylation of bacterial gene promoters is known to influence the promoters affinity for transcription factors, leading to changes in gene expression and epigenetic modifications (23, 41). In this work, we investigated the possible role of the *Wolbachia* MTase as a mediator of gene expression regulation. In absence of promoters, we hypothesise that intragenic DNA methylation could affect gene expression. Indeed, experimental studies revealed that the efficiency of bacterial RNA polymerase is influenced by epigenetic modifications (42–45).

To test our hypothesis, we i) experimentally determined the sequence motif recognised by the wOo DNA MTase; ii) studied the expression data of 55 samples from six published RNA-Seq studies on *Wolbachia*; iii) investigated the presence of regulatory regions upstream the DNA MTase gene in *Wolbachia*.

*In silico* determination of the sequence motif recognized by MTase is challenging (46). This is especially true in this case, considering that this is the first *Wolbachia* DNA MTase enzyme ever characterised. An experimental approach to determining the methylation motif of the enzyme is to express the MTase in an *Escherichia coli* strain depleted for any MTase gene, and then identify the methylated bases and motifs using SMRT PacBio sequencing (46). Surprisingly, wOo MTase was found to be able to methylate both adenine and cytosine, without highly conserved patterns (Figure 1), especially for cytosine. To our knowledge this is the first MTase enzyme described with these peculiar features. Our findings also suggest that the enzyme has a 10-fold greater affinity for cytosine than adenine. This value has been determined on an *E. coli* genome containing ∼50% of AT. Instead, the genome of a *Wolbachia* strain usually contains ∼70% of AT and this partially reduces the MTase affinity bias to 4-fold.

Regarding the analysis of gene expression, we started from the idea that if DNA methylation within the gene affects its expression, we should observe a correlation between gene nucleotide composition and gene expression. Furthermore, the direction (positive or negative) and the strength (the slope of the regression line) of this correlation should change with variation of DNA MTase gene expression. It is well-documented in literature that gene length and codon usage could be correlated to gene expression (46, 47). In particular, highly expressed genes tend to have lower Effective Number of Codons (Nc) and higher Codon Bias Index (CBI). For each of the 55 samples, we computed the correlation between CBI, Nc, gene length, %A, %T, %G, %C and C/A ratio versus the relative gene expression. The result shows correlation between gene nucleotide composition and gene expression in many samples. Interestingly, we found that the slope of the obtained regression lines often correlates with the MTase expression in that sample. In more detail, the expression of genes is positively correlated with their %C in all samples and the correlation slope increases proportionally to the expression of DNA MTase (Figure S2). Regarding %A, the correlation with gene expression was positive (the higher the %A the higher the gene expression) in samples with poorly expressed MTase, while it became negative (the higher the %A the lower the gene expression) in samples with high MTase gene expression (Figure S3).

The positive to negative slope trend observed for %A suggests that methylation of adenine might play an important role in the regulation of gene expression, in particular considering the AT-richness of the *Wolbachia* genome. Lastly, we found that the ratio of %C and %A (C/A ratio) is a good parameter to study *Wolbachia* gene expression: it significantly positively correlates with gene expression in all samples except for those with very low expression of MTase (Figure 2). This strongly supports the hypothesis that the methylations of cytosine and adenine could be the key to understanding the regulation of gene expression in *Wolbachia*. Interestingly, as shown in Figure 4, the genes with low C/A ratio are mainly involved in basal metabolism pathways (such as replication and translation) while those with high C/A ratio are mostly part of pathways involved in stress response (such as the production of energy and ion/molecule transport). Interestingly, among the low or high C/A ratio genes there are, respectively, *fts*Z and *fts*W, both involved in the bacterium cell replication. This suggests that the host could modulate the bacterium replication: a possible key for a successful symbiotic relationship. In this picture, *Wolbachia* could have a stress-related mitochondrion-like role for the host, as previously proposed by Darby and colleagues (2012). Furthermore, it is interesting to note that highly expressed genes have high Codon Bias Index (CBI) (as expected from literature) only for samples with low levels of MTase expression; in those with highly expressed MTase this correlation became not significant (Figure 3). This suggests that the MTase effect on gene expression might have evolved independently from the codon usage adaptation.

The step forward in our analysis was to investigate how the expression of the MTase gene can be regulated. To this aim we analysed a collection of representative *Wolbachia* genomes, screening the 100-bp region upstream of the MTase gene for the presence of regulatory sequences. This analysis highlighted the presence of CtrA binding site and TATA box sequences highly conserved among *Wolbachia* strains. The CtrA binding site is a sequence bounded by the CtrA response regulator protein, which is part of the two component regulatory system Ccka/CtrA. This system is composed of Ccka, a membrane sensor histidine kinase, and CtrA, an intracellular effector protein. When triggered by an external stimulus, the Ccka protein activates CtrA, which in turn binds to the CtrA binding site initiating the expression of the downstream genes. While the signalling mechanisms behind the activation of Ccka are not fully understood, the downstream response regulator, CtrA, is a well-described master regulator with transcription factor activity, conserved in *Alphaproteobacteria* (*48*). The presence of a highly conserved CtrA binding site upstream the MTase gene locus, strongly suggests that the CcKA/CtrA signal transduction pathway could have a role in the regulation of the expression of this gene. Indeed, the host could modulate the expression of the *Wolbachia* MTase gene by stimulating the Ccka receptor on the membrane on the bacterial cell. Furthermore, the presence of a conserved CtrA binding site upstream of only two genes suggests that this could be a highly regulated mechanism subjected to a strong selective pressure.

Surprisingly, we also found highly conserved TATA boxes (i.e. eukaryotic regulatory sequences) upstream of the MTase gene. In eukaryotes, TATA boxes tend to be placed upstream of stress-responsive genes, whose expression must rapidly and variably be tuned in response to specific environmental conditions and changing physiological needs (49, 50) . The TATA box is bound by a specific protein, the TATA box binding protein (TBP), which then recruits transcription factors. *Wolbachia* does not encode TBP proteins, so the functionality of these TATA box sequences can only be hypothesised, e.g. assuming that TBPs are supplied by the host. Interestingly, we found 12 *Wolbachia* genes having highly conserved TATA boxes. These genes include *ctr*A response regulator (see above), genes involved in energy production and a component of the Type IV secretion system.

To explain how the host could regulate gene expression in *Wolbachia*, we propose the following model: i) specific molecules in the host cells activate the Ccka membrane histidine kinase; ii) Ccka phosphorylates the intracellular CtrA response regulator; iii) the phosphorylated CtrA binds to the CtrA binding site upstream of the MTase gene inducing its expression in *Wolbachia*; iv) the MTase enzyme methylates adenine and cytosine of the *Wolbachia* genome; v) the expression of several *Wolbachia* genes changes on the basis of their nucleotide content, particularly enhancing the transcription of genes with a high ratio of cytosine / adenine. The model is graphically summarised in Figure 5.

**Figure 5.**
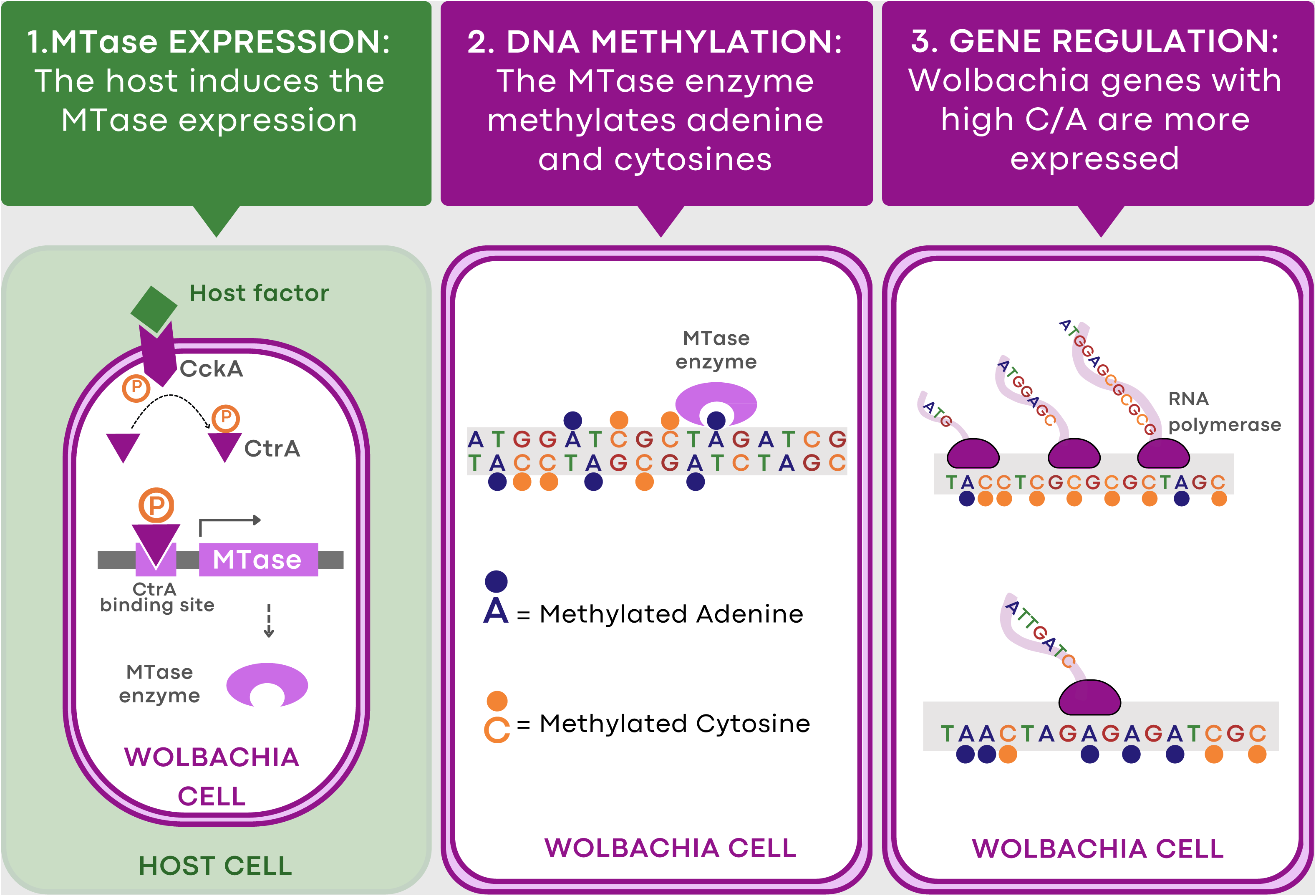
Graphical representation of the mechanism proposed for the regulation of the *Wolbachia* gene expression The figure summarises the model here proposed for the *Wolbachia pipientis* gene expression regulation. *Wolbachia* is an intracellular bacterium able to colonise the cytoplasm of host cells. In the figure the bacterium, represented with violet membranes and white cytoplasm, is surrounded by the host cytoplasm, coloured in green. At step 1 (on the left): the host activates the Ccka sensor histidine kinase (in violet) placed on the *Wolbachia* membrane (here we represented this activation as mediated by a molecular factor produced by the host, but other mechanisms are possible), then the Ccka phosphorilates the intracellular transcription regulator CtrA which, induces the transcription of the *Wolbachia* DNA *methyltransferase* gene (MTase) binding the CtrA binding site upstream it. At step 2 (in the middle): the MTase enzyme methylates the adenines and cytosines of the *Wolbachia* DNA and methyl groups are represented with dots coloured as the bases. At step 3 (on the right): RNA polymerases are represented by pink icons from which mRNA molecules originate. In highly expressed genes more RNA polymerases bind the gene simultaneously. The greater is the cytosines / adenines ratio in a gene the more that gene is expressed.

Moreover, the presence of a conserved TATA box upstream the *ctr*A gene could suggest a possible role for the host in the regulation of *ctr*A gene expression.

While the hypothesised mechanism may not lead to highly precise regulation of gene expression, it could be a rough yet functional mechanism, sufficient for the establishment of a successful intracellular symbiosis. In the future, it would be interesting to investigate whether similar mechanisms are present in other endosymbiont or intracellular organisms.

Despite the results presented in this work are far from being definitive, we hope to have opened a new way in understanding the symbiotic relationship between *Wolbachia* and its hosts. In our opinion, this is a magnificent example of how evolution works, recycling metabolic pieces for similar aims, sometimes reaching surprisingly stable *equilibria*.

## Acknowledgments

F.C. would like to thank his mentor, Claudio Bandi, for introducing him to the world of *Wolbachia* and for his helpful comments to this paper.

