## Supplemental figures for "Evidence that host-mediated epigenetic modifications regulate gene expression in *Wolbachia pipientis*"

**Eukaryotic hosts could regulate the gene expression of the endosymbiont bacterium *Wolbachia pipientis***

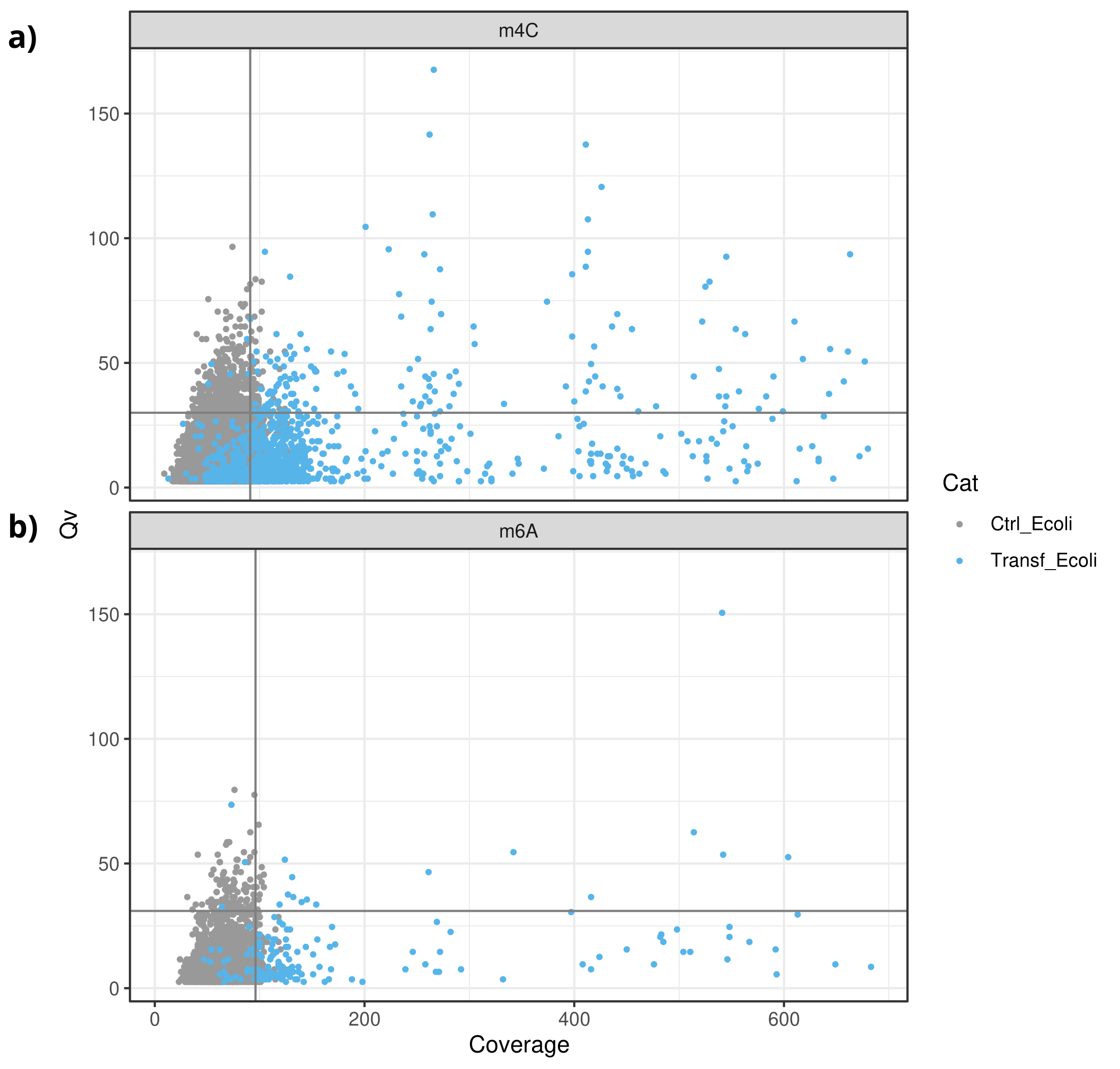

**Figure S1. Scatter plot of Qv and Coverage of methylated motifs**

The Qv and coverage values of all the putative methylated motifs obtained from the control and the transformed Stellar Escherichia coli (dam-/dcm-) strain were studied to determine the threshold for the calling of methylated positions. Azure points refers to methylation sites on the transformed E. coli strain, while gray on the control strain. Horizontal and vertical lines identify respectively the 95 percentile of Qv and coverage values for the control E. coli strain.

The Qv vs coverage scatter plot relative to m4C methylations of cytosines is reported in plot (a), for m6A methylation of adenines in plot (b).

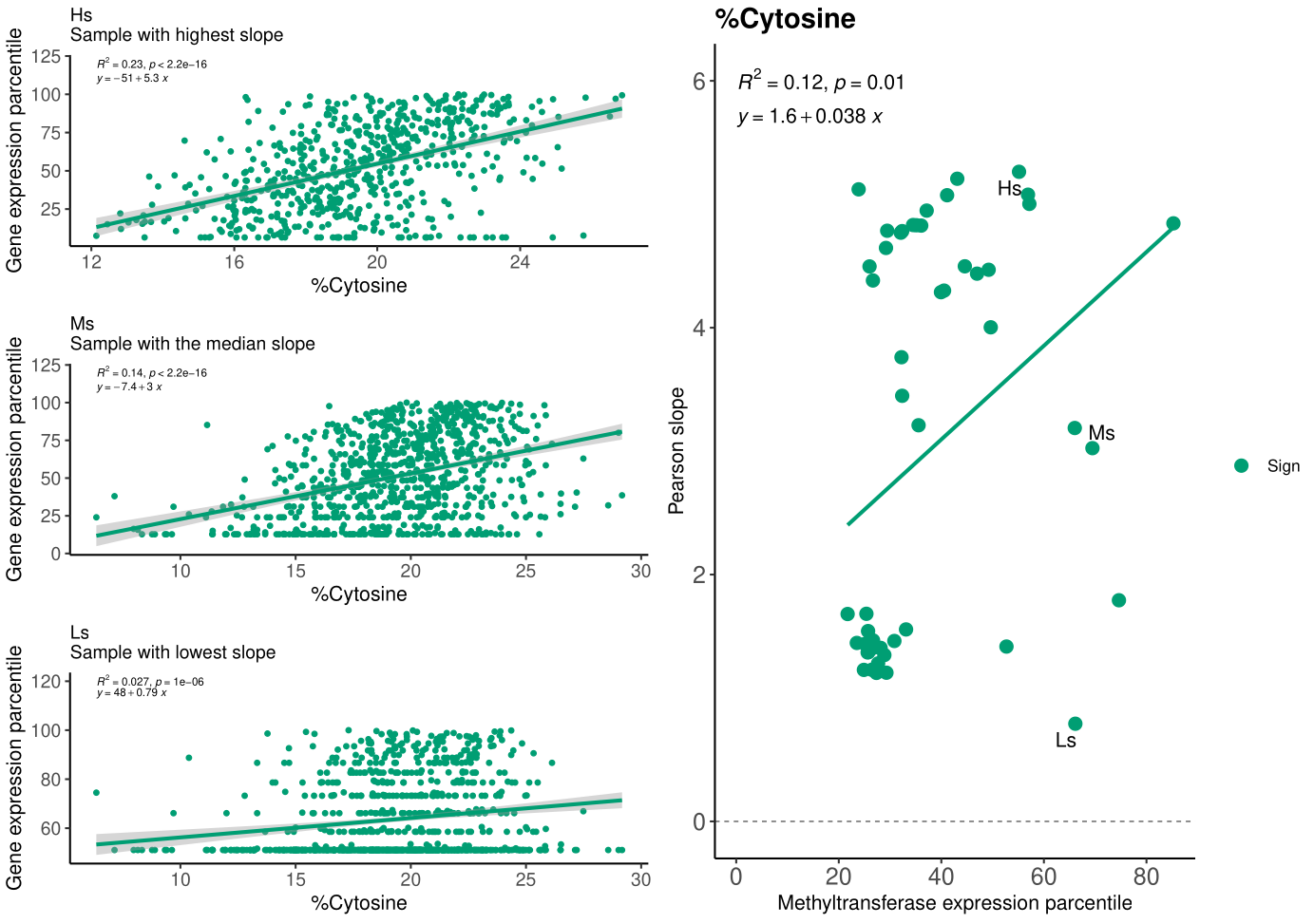

**Figure S2. Correlation between the expression of genes and their percentage of cytosine**

For each of the 55 samples included in the study, the correlation between the expression of the genes and the relative percentage of cytosine (%C) was investigated. The scatter plots of %C vs gene expression of three exemplificative samples selected from the 55 samples are reported: a) the correlation plot with highest slope (Hs), b) the correlation plot with the medium slope (Ms) and c) the correlation plot with the lowest slope (Ls). In each of these three plots, colour is green whether the correlation is significant (p value < 0.05) and grey if it is not, the regression line is coloured in dark grey and the statistics are reported on the chart. In figure d, the correlation plot between the slopes obtained for each of the 55 samples, and the relative expression of the DNA methyltransferase gene is reported. Points relative to significant correlations (p value < 0.05) are in green while non significant in grey. The regression line is in green and the statistics are reported on the chart.

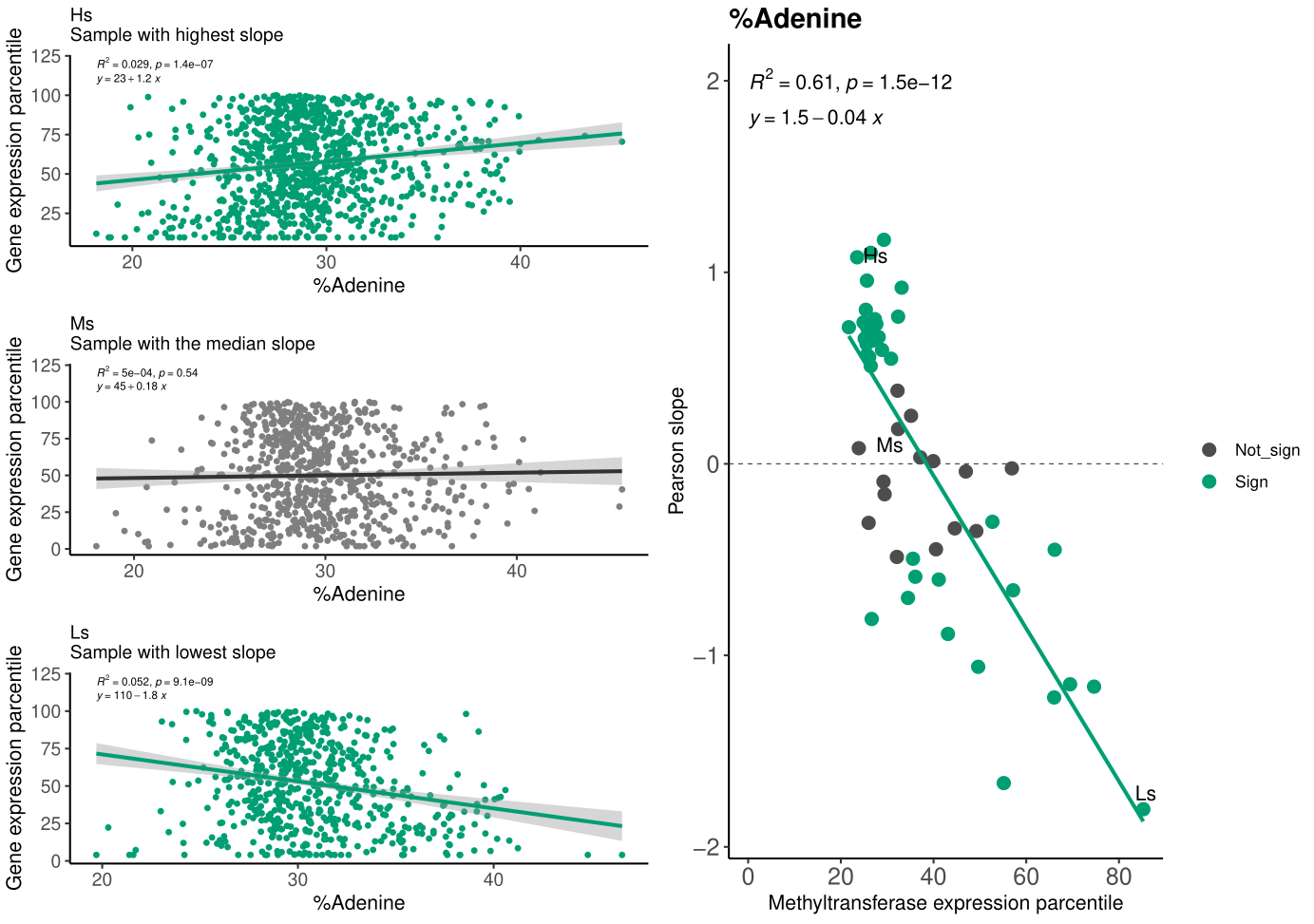

**Figure S3. Correlation between the expression of genes and their percentage of adenine**

For each of the 55 samples included in the study, the correlation between the expression of the genes and the relative percentage of adenine (%A) was investigated. The scatter plots of %A vs gene expression of three exemplificative samples selected from the 55 samples are reported: a) the correlation plot with highest slope (Hs), b) the correlation plot with the medium slope (Ms) and c) the correlation plot with the lowest slope (Ls). In each of these three plots, colour is green whether the correlation is significant (p value < 0.05) and grey if it is not, the regression line is coloured in dark grey and the statistics are reported on the chart. In figure d, the correlation plot between the slopes obtained for each of the 55 samples, and the relative expression of the DNA methyltransferase gene is reported. Points relative to significant correlations (p value < 0.05) are in green while non significant in grey. The regression line is in green and the statistics are reported on the chart.

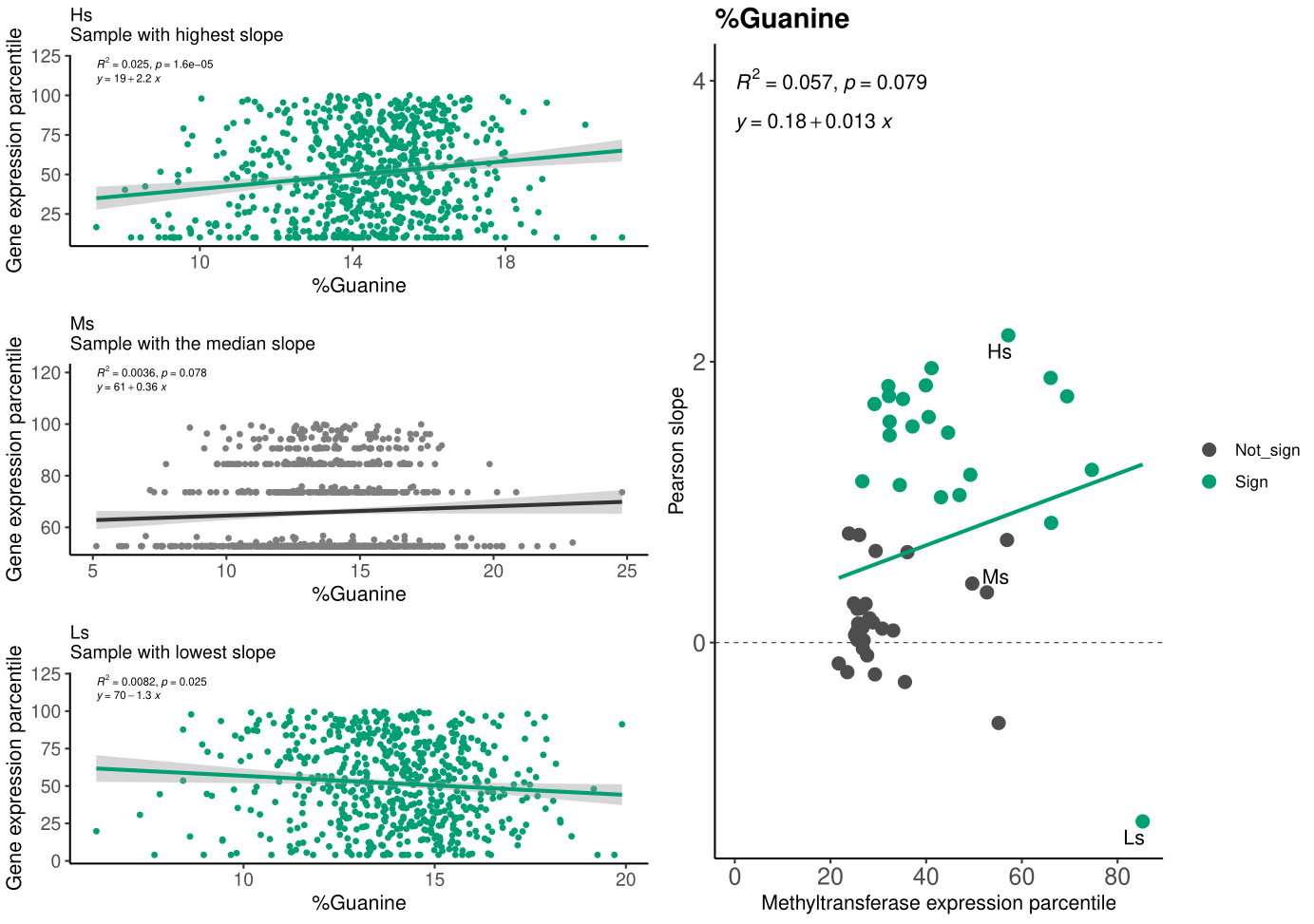

**Figure S6. Correlation between the expression of genes and their percentage of guanine**

For each of the 55 samples included in the study, the correlation between the expression of the genes and the relative percentage of guanine (%G) was investigated. The scatter plots of %G vs gene expression of three exemplificative samples selected from the 55 samples are reported: a) the correlation plot with highest slope (Hs), b) the correlation plot with the medium slope (Ms) and c) the correlation plot with the lowest slope (Ls). In each of these three plots, colour is green whether the correlation is significant (p value < 0.05) and grey if it is not, the regression line is coloured in dark grey and the statistics are reported on the chart. In figure d, the correlation plot between the slopes obtained for each of the 55 samples, and the relative expression of the DNA methyltransferase gene is reported. Points relative to significant correlations (p value < 0.05) are in green while non significant in grey. The regression line is in green and the statistics are reported on the chart.

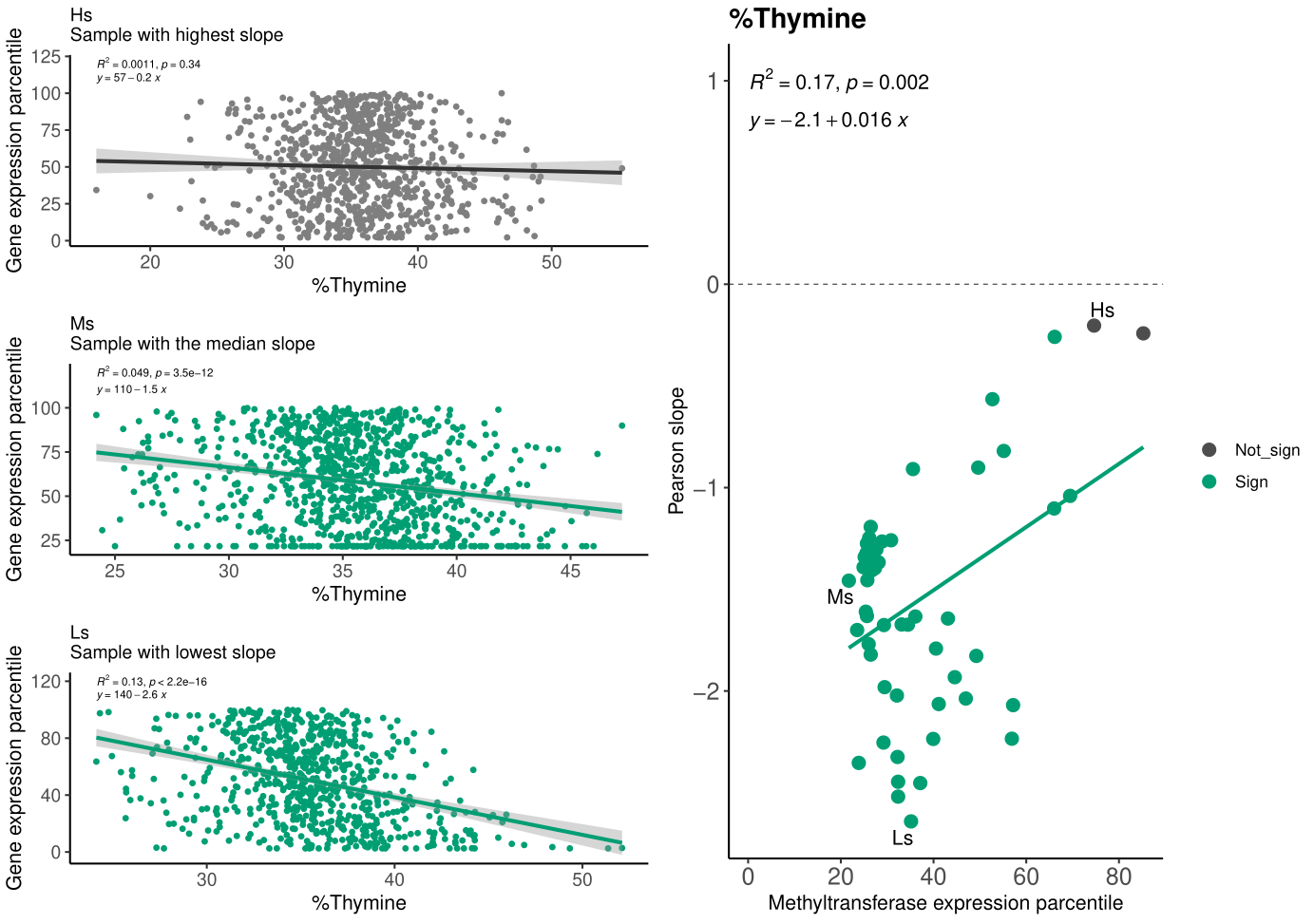

**Figure S5. Correlation between the expression of genes and their percentage of thymine**

For each of the 55 samples included in the study, the correlation between the expression of the genes and the relative percentage of thymine (%T) was investigated. The scatter plots of %T vs gene expression of three exemplificative samples selected from the 55 samples are reported: a) the correlation plot with highest slope (Hs), b) the correlation plot with the medium slope (Ms) and c) the correlation plot with the lowest slope (Ls). In each of these three plots, colour is green whether the correlation is significant (p value < 0.05) and grey if it is not, the regression line is coloured in dark grey and the statistics are reported on the chart. In figure d, the correlation plot between the slopes obtained for each of the 55 samples, and the relative expression of the DNA methyltransferase gene is reported. Points relative to significant correlations (p value < 0.05) are in green while non significant in grey. The regression line is in green and the statistics are reported on the chart.

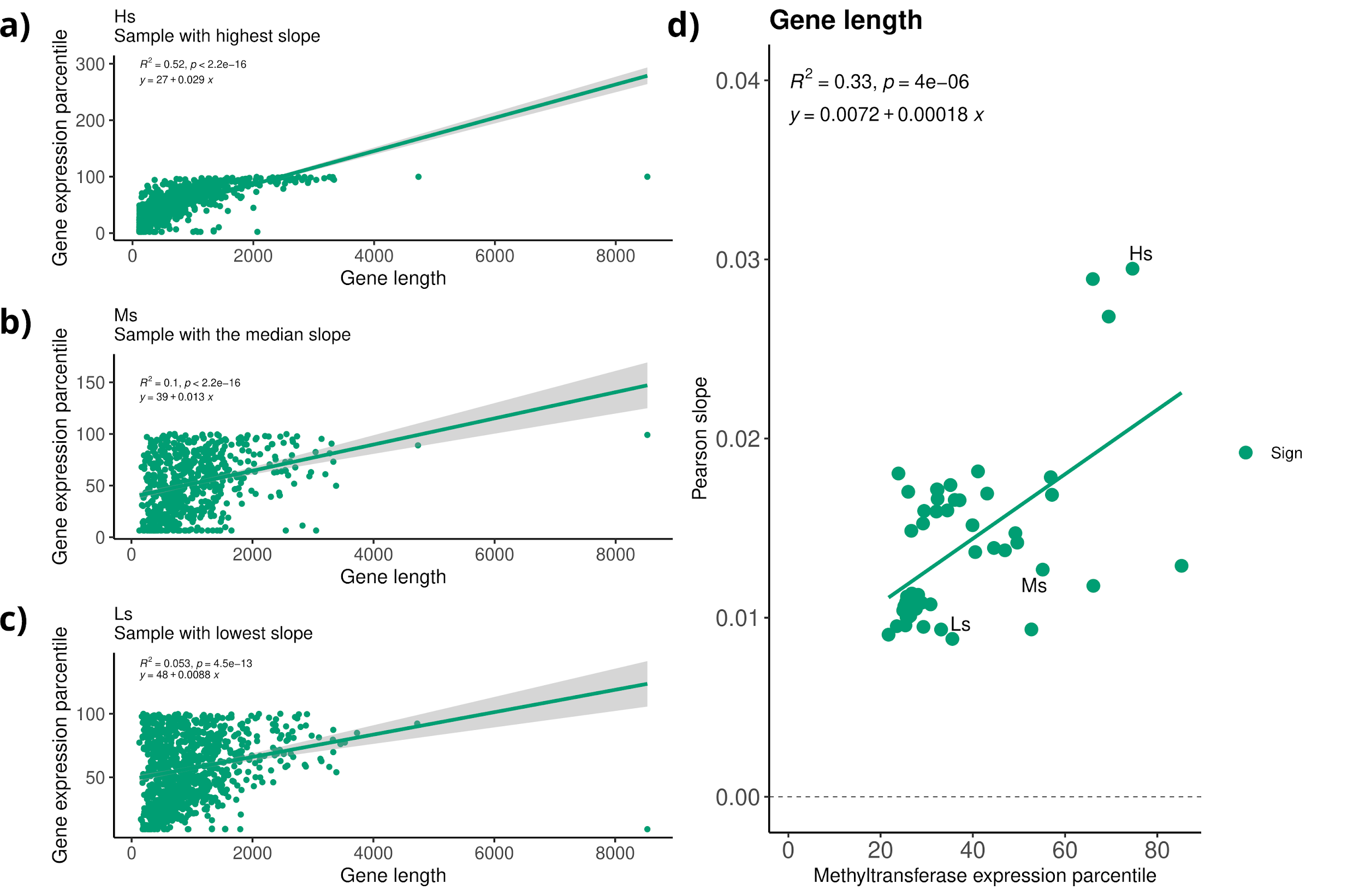

**Figure S6. Correlation between the expression of genes and their length**

For each of the 55 samples included in the study, the correlation between the expression of the genes and the relative gene length was investigated. The scatter plots of gene length vs gene expression of three exemplificative samples selected from the 55 samples are reported: a) the correlation plot with highest slope (Hs), b) the correlation plot with the medium slope (Ms) and c) the correlation plot with the lowest slope (Ls). In each of these three plots, colour is green whether the correlation is significant (p value < 0.05) and grey if it is not, the regression line is coloured in dark grey and the statistics are reported on the chart. In figure d, the correlation plot between the slopes obtained for each of the 55 samples, and the relative expression of the DNA methyltransferase gene is reported. Points relative to significant correlations (p value < 0.05) are in green while non significant in grey. The regression line is in green and the statistics are reported on the chart.

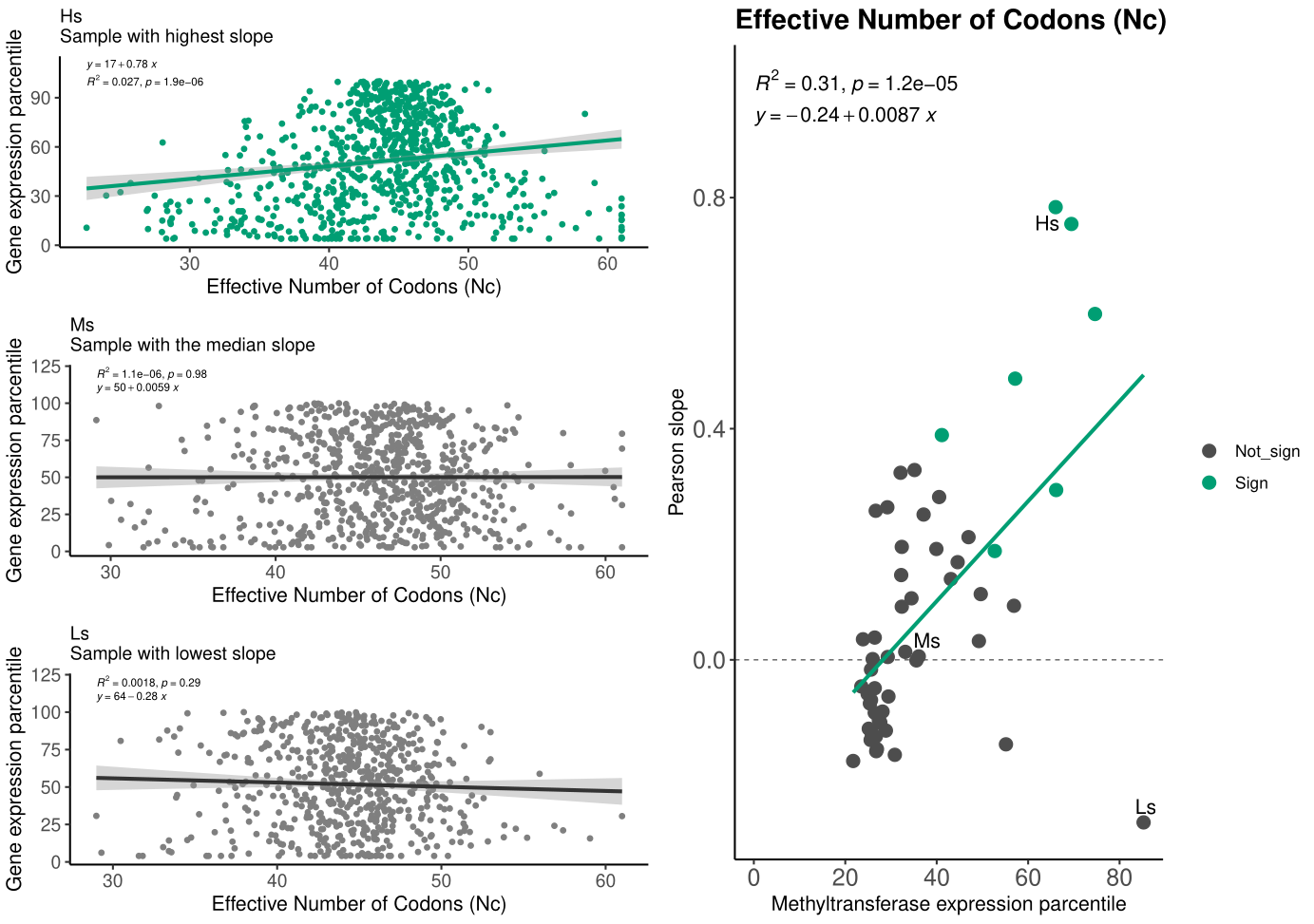

**Figure S7. Correlation between the expression of genes and their Effective number of codons (Nc)**

For each of the 55 samples included in the study, the correlation between the expression of the genes and the relative Effective number of codons (Nc) was investigated. The scatter plots of Nc vs gene expression of three exemplificative samples selected from the 55 samples are reported: a) the correlation plot with highest slope (Hs), b) the correlation plot with the medium slope (Ms) and c) the correlation plot with the lowest slope (Ls). In each of these three plots, colour is green whether the correlation is significant (p value < 0.05) and grey if it is not, the regression line is coloured in dark grey and the statistics are reported on the chart. In figure d, the correlation plot between the slopes obtained for each of the 55 samples, and the relative expression of the DNA methyltransferase gene is reported. Points relative to significant correlations (p value < 0.05) are in green while non significant in grey. The regression line is in green and the statistics are reported on the chart.

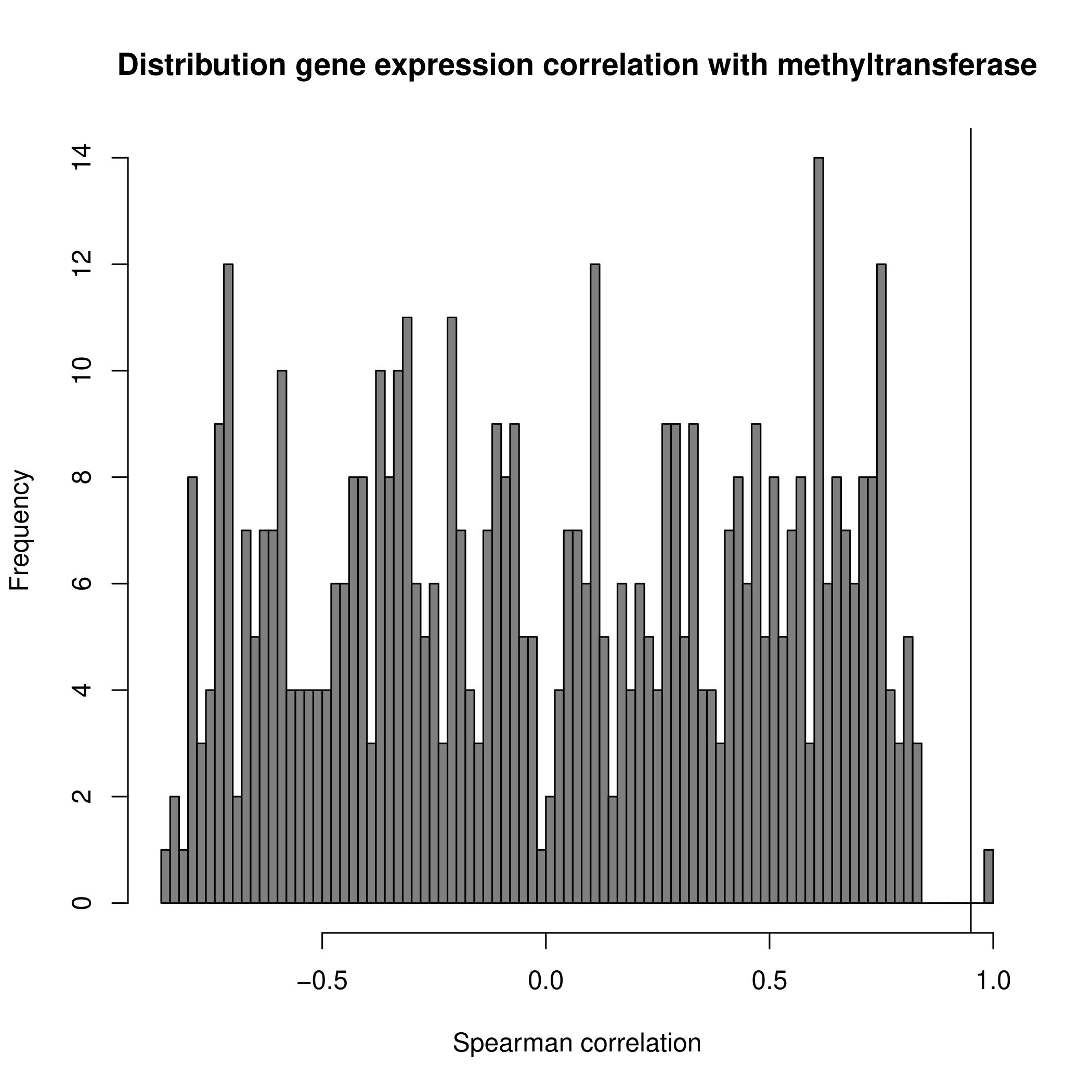
**Figure S8. Distribution of the expression distances from the DNA methyltransferase**

The gene expression of all the core genes present among all the 55 samples included in the study were compared by Spearman co-correlation. The distribution of the distances from the DNA methyltransferase gene is reported. The vertical line identifies the 0.9 distance value. The correlation value 1 is due to the gene against itself.

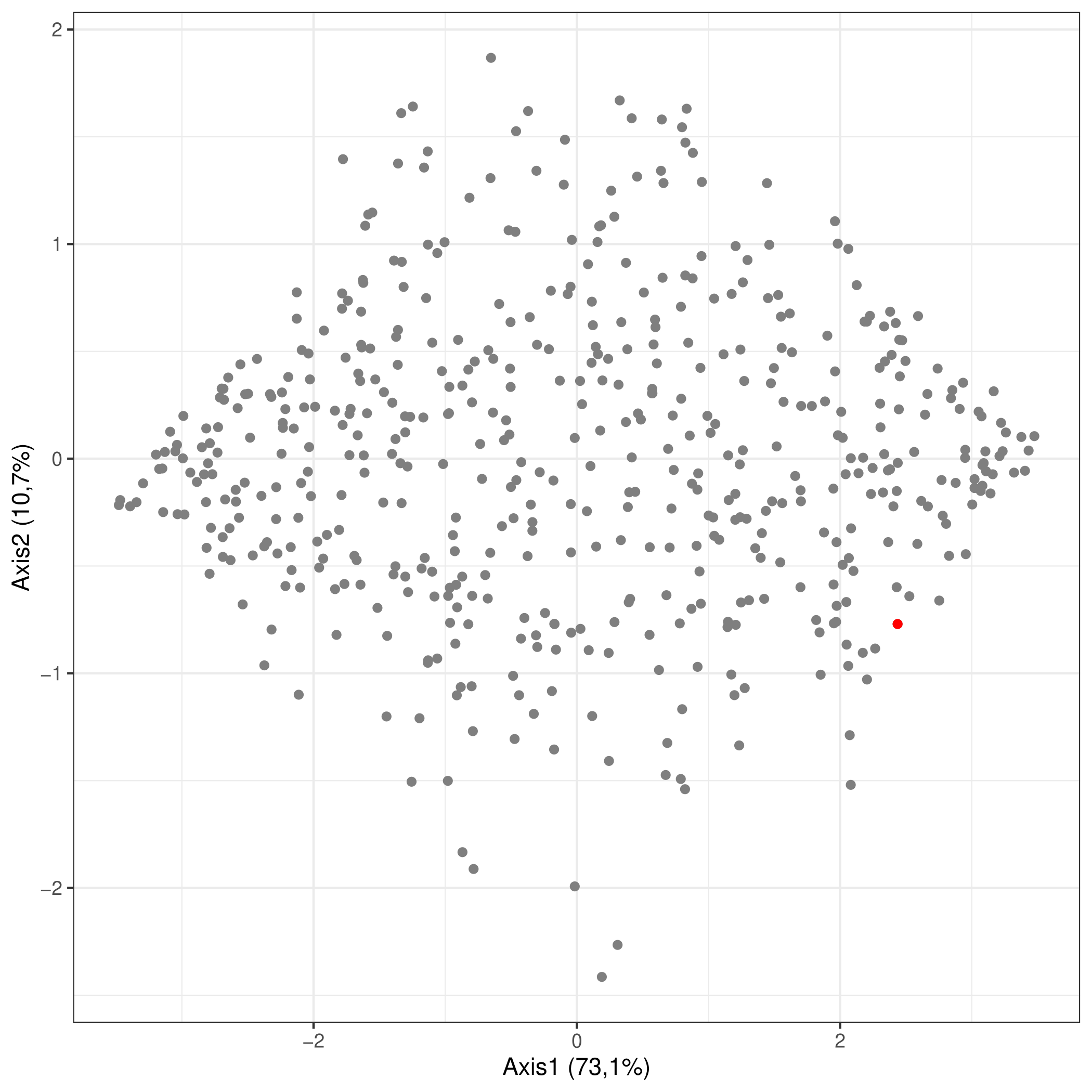
**Figure S9. Principal Component Analysis (PCA) on the expression of all core genes**

The gene expression of all the core genes present among all the 55 samples included in the study were subjected to Principal Component Analysis (PCA) for the detection of gene, or cluster of gene, with an expression pattern similar to DNA methyltransferase (in red in the plot).

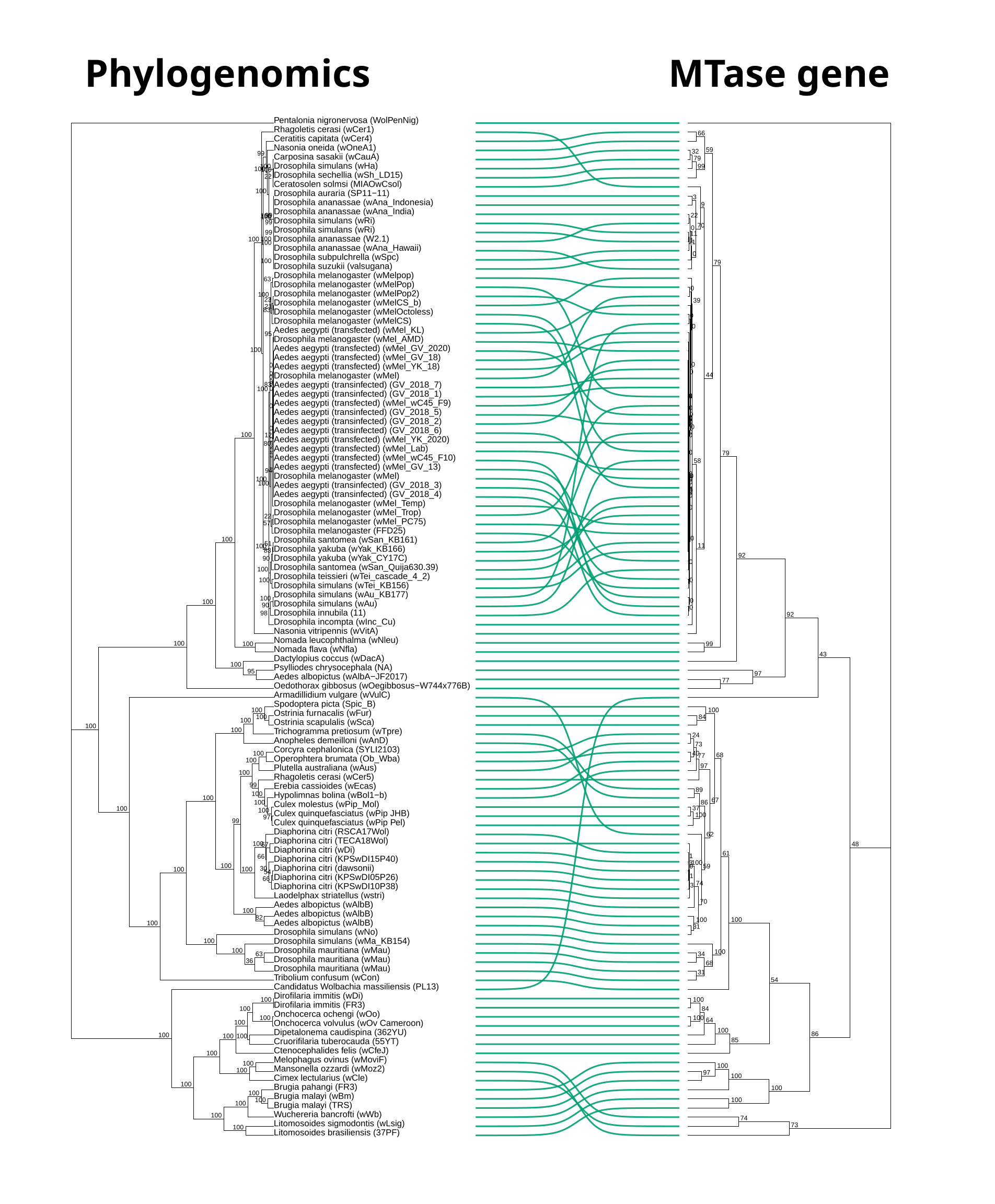
**Figure S10. Comparison of *Wolbachia pipientis* vs MTase phylogenetic trees**

The Maximum Likelihood (ML) trees obtained from: on the left, the nucleotide concatenate of the single copy core genes of 112 representative Wolbachia genomes; on the right, the nucleotide sequences of the 112 MTase gene retrieved from the same genomic dataset. Green lines connect the corresponding strains on the trees and bootstrap support values are reported in the trees.

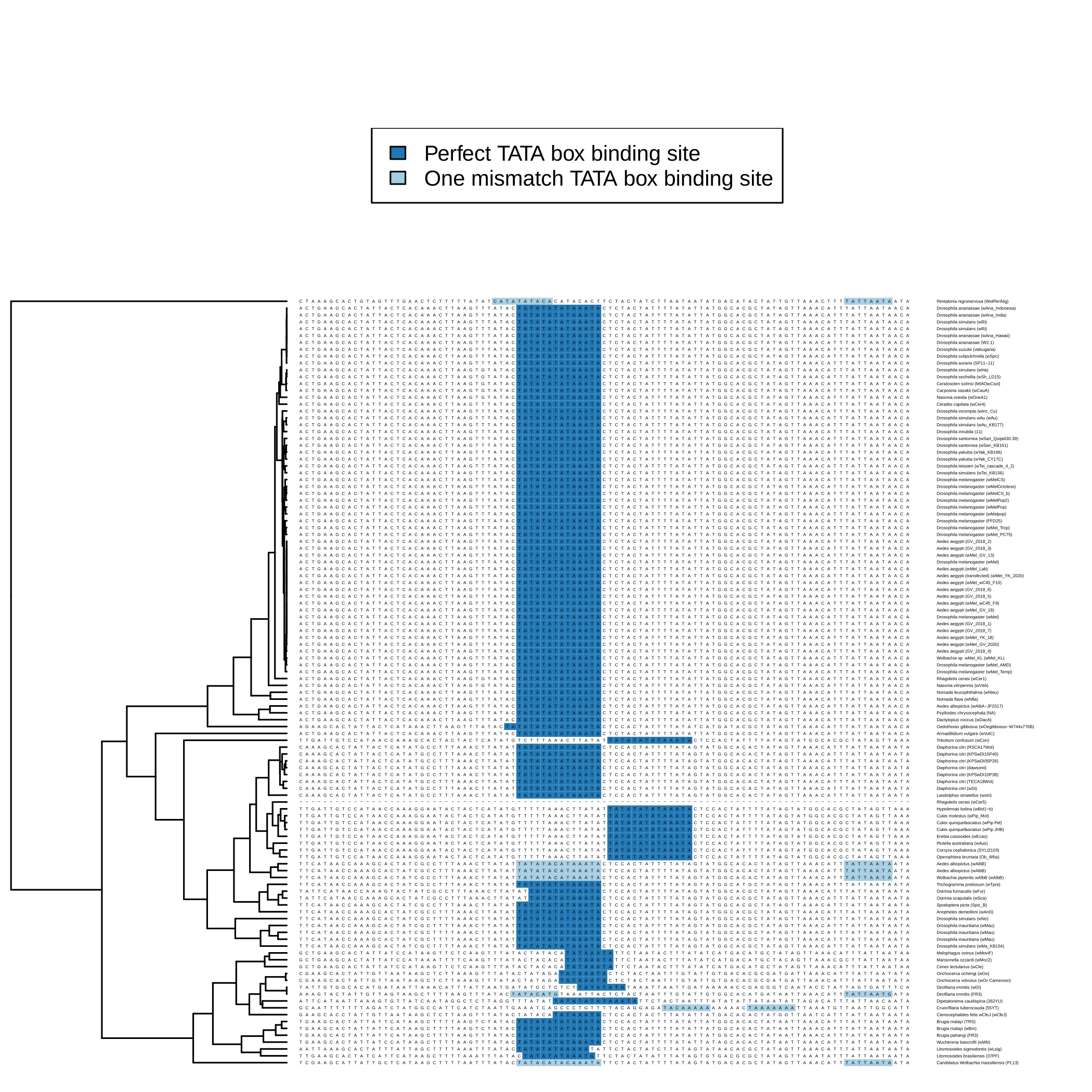

**Figure S11. TATA boxes in the 100 bp regions upstream the DNA methyltransferase gene**

The 100 bp regions upstream of the DNA methyltransferase gene (MTase) on 111 representative Wolbachia strains are shown. On the left, the Maximum Likelihood (ML) phylogenetic analysis obtained from the concatenate of the single copy core genes. On the right the perfect TATA boxes in blue and one mismatched TATA boxes in azure. One sequence (GCF_018454445.1, Wolbachia endosymbiont of Rhagoletis cerasi - wCer5) is composed only by gap because the MTase gene was placed on the extreme of the contig and it was not possible to retrieve the upstream region.
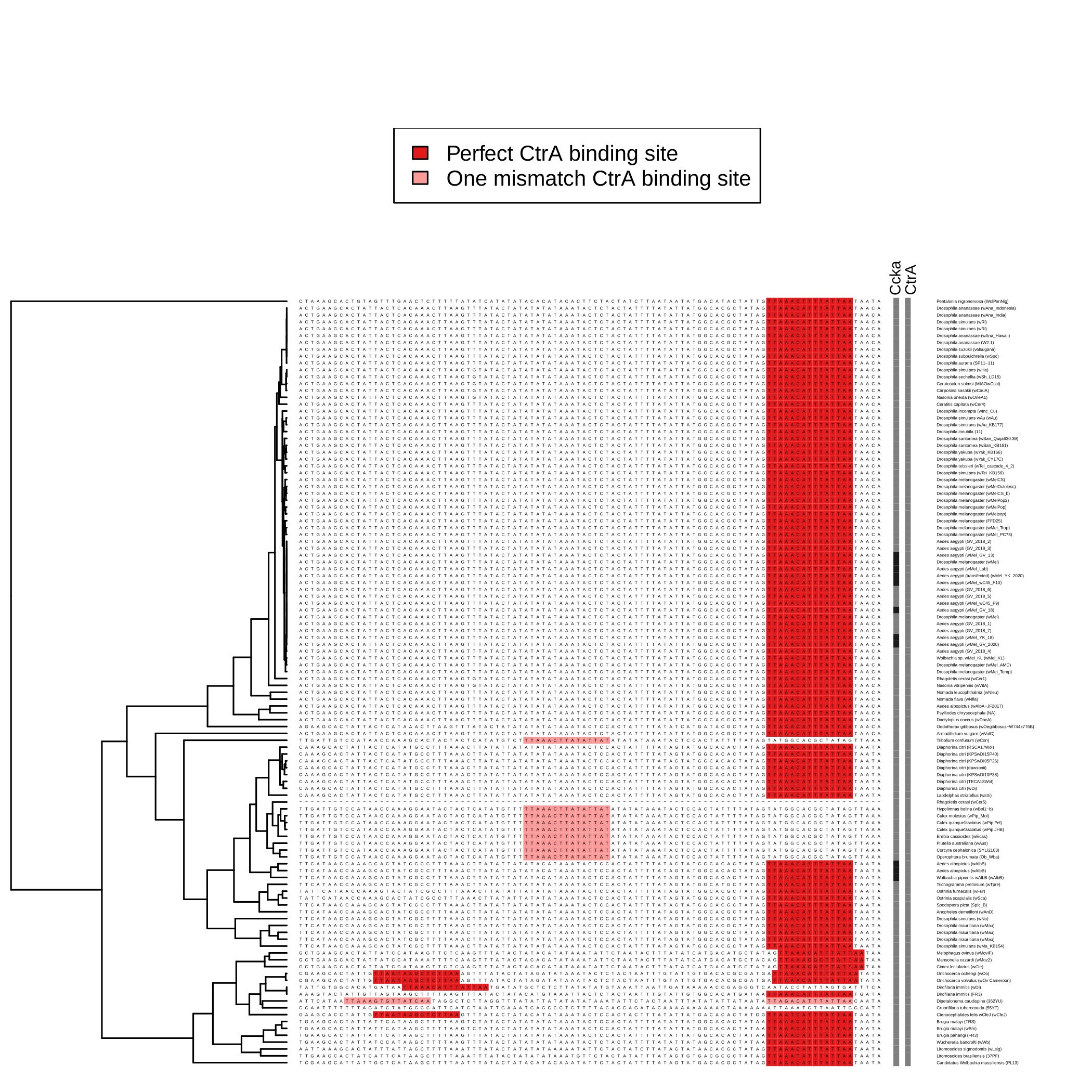

**Figure S12. CtrA boxes in the 100 bp regions upstream the DNA methyltransferase gene**

The 100 bp regions upstream of the DNA methyltransferase gene (MTase) on 111 representative Wolbachia strains are shown. On the left, the Maximum Likelihood (ML) phylogenetic analysis obtained from the concatenate of the single copy core genes. On the right are the perfect CtrA boxes in red and one mismatched TATA boxes in light red. One sequence (GCF_018454445.1, Wolbachia endosymbiont of Rhagoletis cerasi - wCer5) is composed only by gap because the MTase gene was placed on the extreme of the contig and it was not possible to retrieve the upstream region.

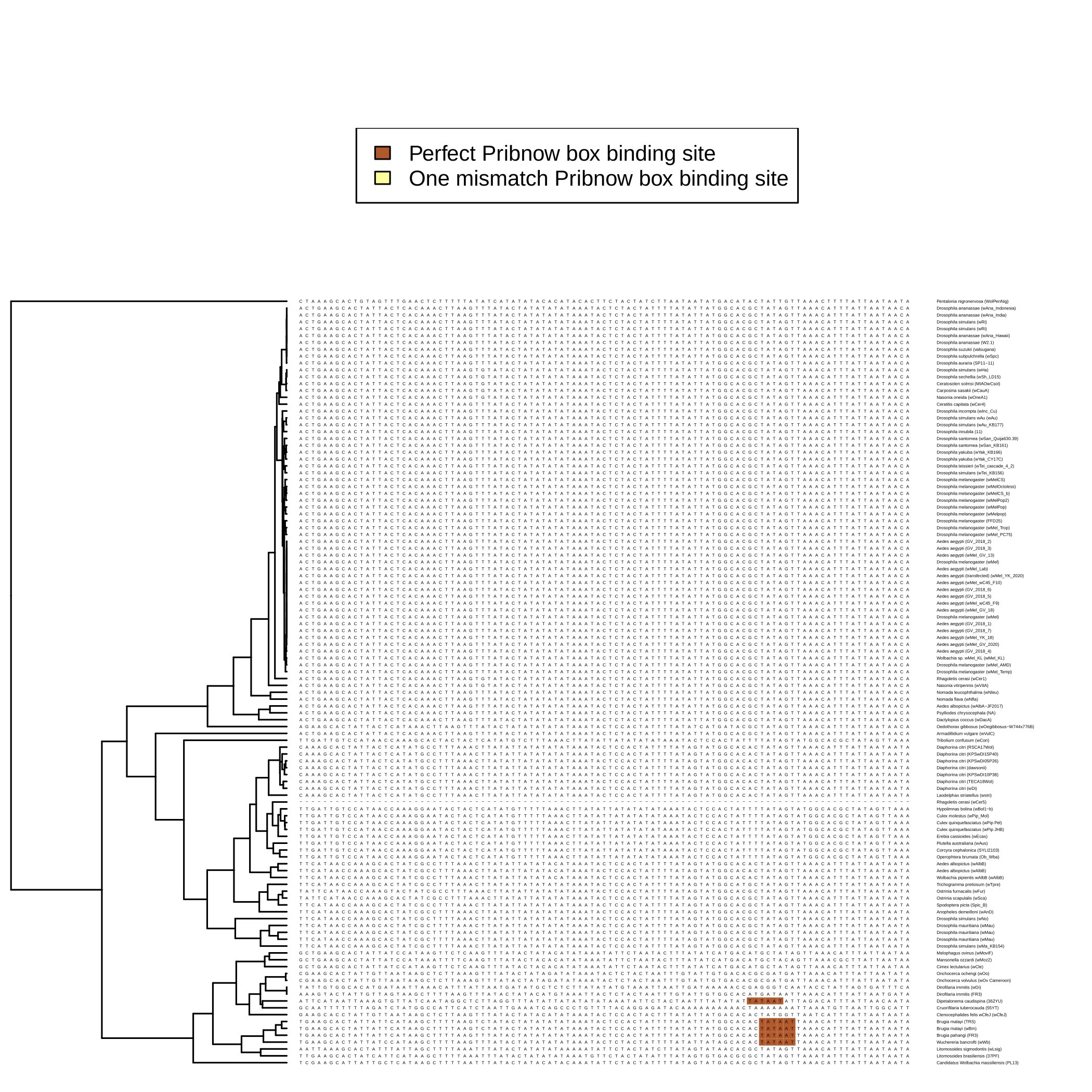

**Figure S13. Pribnow boxes in the 100 bp regions upstream the DNA methyltransferase gene**

The 100 bp regions upstream of the DNA methyltransferase gene (MTase) on 111 representative Wolbachia strains are shown. On the left, the Maximum Likelihood (ML) phylogenetic analysis obtained from the concatenate of the single copy core genes. On the right the perfect Pribnow boxes in brown. One sequence (GCF_018454445.1, Wolbachia endosymbiont of Rhagoletis cerasi - wCer5) is composed only by gap because the MTase gene was placed on the extreme of the contig and it was not possible to retrieve the upstream region.

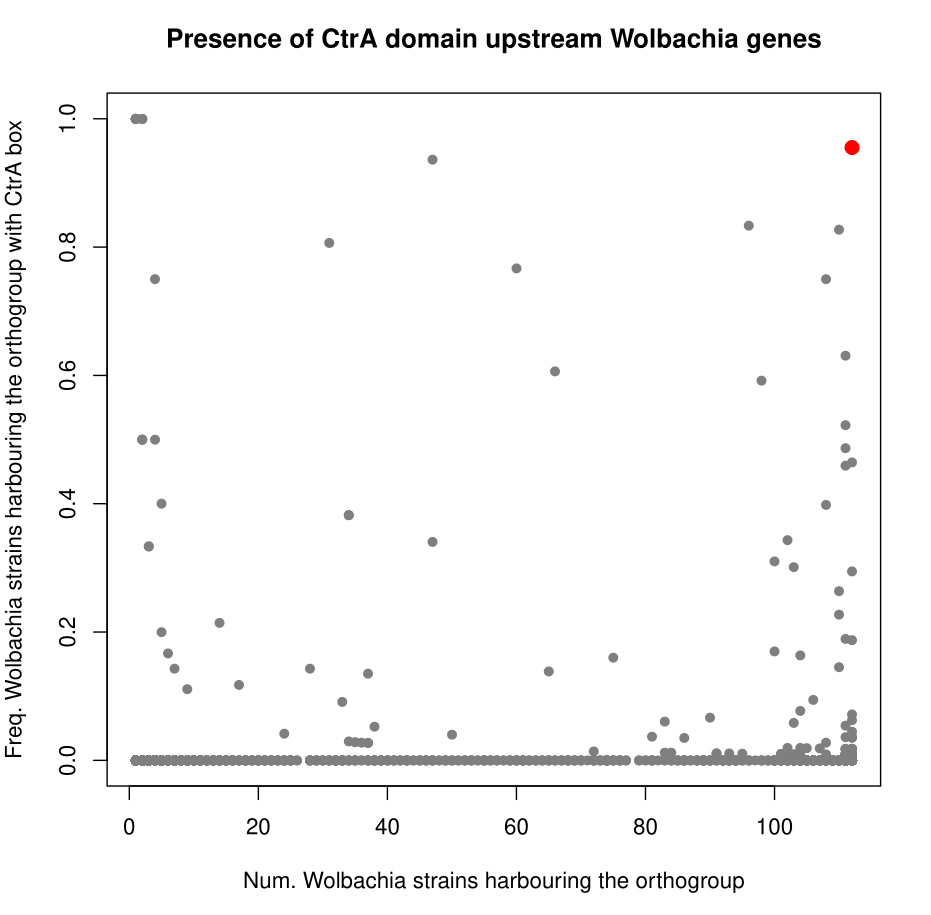

**Figure S14. Scatter plot between ortholog gene and CtrA binding domains occurrence**

Scatter plot in which each point is one of the ortholog genes found among the 112 Wolbachia genomes included in the study. On the x-axis, the number of Wolbachia genomes harbouring the ortholog gene and on the y-axis the frequency of sequences preceded by CtrA binding domain. The red dot correspond to the MTase gene.

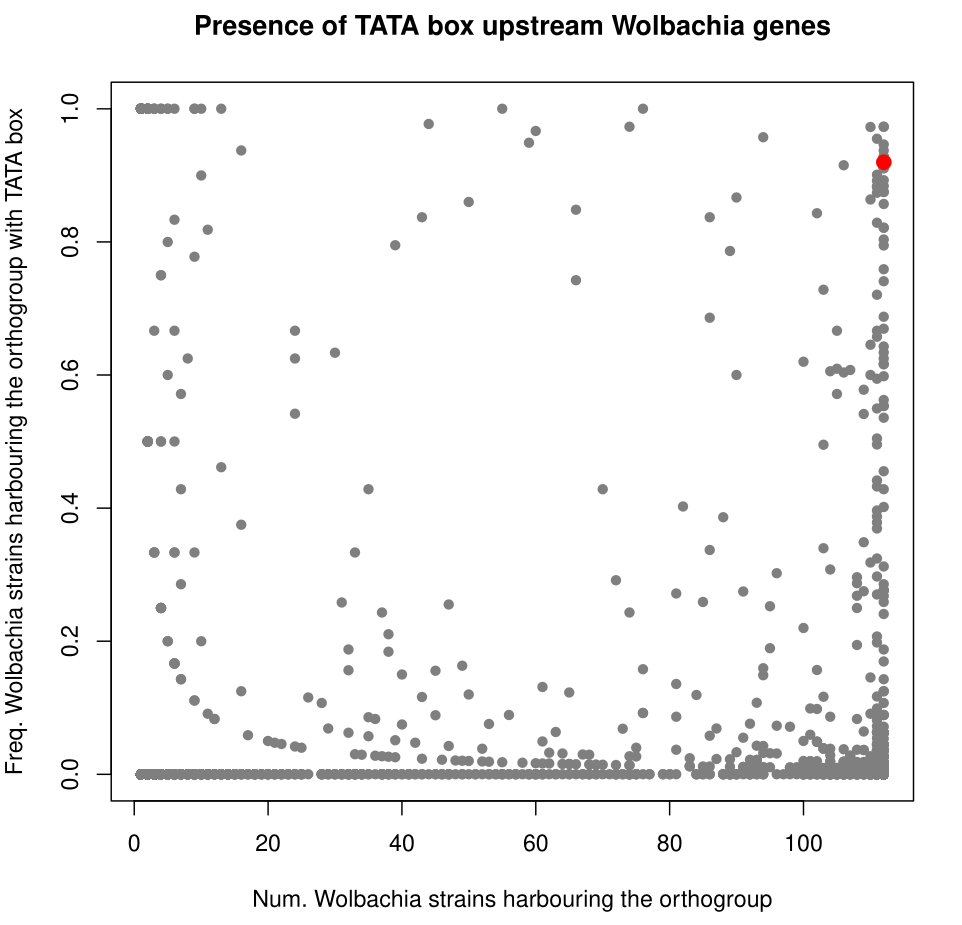

**Figure S15. Scatter plot between ortholog gene and TATA box occurrence**

Scatter plot in which each point is one of the ortholog genes found among the 112 Wolbachia genomes included in the study. On the x-axis, the number of Wolbachia genomes harbouring the ortholog gene and on the y-axis the frequency of sequences preceded by TATA box. The red dot correspond to the MTase gene.
